# Bridging model and experiment in systems neuroscience with Cleo: the Closed-Loop, Electrophysiology, and Optophysiology simulation testbed

**DOI:** 10.1101/2023.01.27.525963

**Authors:** Kyle A. Johnsen, Nathanael A. Cruzado, Zachary C. Menard, Adam A. Willats, Adam S. Charles, Jeffrey E. Markowitz, Christopher J. Rozell

## Abstract

Systems neuroscience has experienced an explosion of new tools for reading and writing neural activity, enabling exciting new experiments (e.g., all-optical interrogation, closed-loop control) for interrogating neural circuits. Unfortunately, these advances have drastically increased the complexity of designing experiments, with ad hoc decisions often resulting in suboptimal or even failed experiments. Bridging model and experiment via simulation can help solve this problem, leveraging advances in computational models to provide a low-cost testbed for experiment design, model validation, and methods engineering. Specifically, we require an integrated approach that incorporates simulation of the experimental interface into computational models, but no existing tool integrates optogenetics, two-photon calcium imaging, electrode recording, and flexible closed-loop processing with neural population simulations. To address this need, we have developed Cleo: the Closed-Loop, Electrophysiology, and Optophysiology experiment simulation testbed. Cleo is a Python package enabling injection of virtual recording and stimulation devices as well as closed-loop control with realistic latency into a Brian spiking neural network model. Notably, it is the only publicly available tool to date simulating two-photon and multi-opsin/wavelength optogenetics. To facilitate adoption and extension by the community, Cleo is open-source, modular, tested, and documented, and can export results to various data formats. Here we describe the design and features of Cleo, evaluate output of individual components and integrated experiments, and demonstrate its utility for advancing optogenetic techniques in prospective experiments using previously published systems neuroscience models.

## 1. Introduction

Systems neuroscience is currently undergoing a revolution fueled by advances in neural manipulation (Fenno et al., 2011; Adesnik and Abdeladim, 2021), measurement (Steinmetz et al., 2021; Gutruf and Rogers, 2018; Knöpfel and Song, 2019; Svoboda and Yasuda, 2006), and analysis (Berman et al., 2014; Mathis et al., 2018; Cunningham and Yu, 2014) methods. These have yielded unprecedented datasets, insights into network activity, and novel paradigms such as direct closed-loop control of neural activity (Grosenick et al., 2015; Potter et al., 2014; Acharya et al., 2022; Bolus et al., 2018, 2021; Zhang et al., 2018; Bergs et al., 2023; Krook-Magnuson et al., 2013; Witt et al., 2013; Dutta et al., 2019; Shang et al., 2024). While exciting, this explosion in the sophistication and quantity of techniques has led to missed opportunities, untapped potential, and even failed experiments (Taniguchi et al., 2024) as it has become difficult to adequately select from an ever-growing catalog of tools via mental models or ad hoc design processes alone.

One solution lies in low-cost simulations that prototype or validate experimental parameters and goals before expensive *in vivo* implementation. This approach requires an integrated model of measurement and manipulation tools together with the brain (see Fig. 1a), forming a digital twin of the experiment. Integrated models are crucial for complex tools whose limitations, idiosyncrasies, and interactions can determine experiment success or failure. For example, closed-loop stimulation involves real-time interaction with the neural system, making experimental design more difficult.

**Figure 1:**
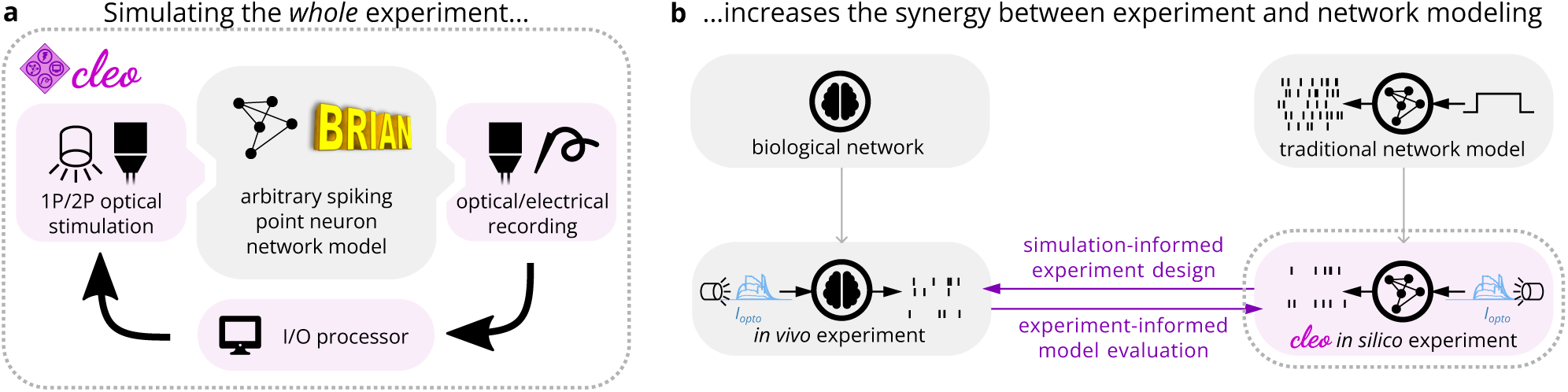
Cleo enables simulation of complex systems neuroscience experiments, augmenting the synergy between model and experiment. (A) Cleo incorporates into a network model components (pink shading) rarely simulated in computational neuroscience models. It does so by wrapping a Brian network model, injecting virtual stimulation and recording devices, and interfacing with the network through a simulated “I/O processor” to control stimulation in an optionally closed-loop and/or delayed fashion. (B) As an experiment simulation testbed, Cleo allows experiment and model to better inform each other. By simulating the measurement and manipulation of the underlying neural activity, Cleo produces simulation results that are more directly comparable to *in vivo* experiments.

Beyond facilitating experiment design, integrated experiment modeling can be viewed more generally as bridging computational and experimental neuroscience, providing additional benefits as the two fields better inform each other. One benefit is enhanced model development through direct comparison to experimental data. For example, computational models can be validated against data from optogenetics/electrophysiology experiments with greater confidence by simulating dynamic photocurrents and noisy spike detection rather than by simply injecting synthetic currents and perfectly recording every spike (see Fig. 1b). Engineering new proteins or analytic methods can also benefit from a testbed that includes algorithms, opsins, and sensors. The NAOMi calcium imaging simulator (Song et al., 2021), for example, has been a valuable resource in validating and tuning segmentation, denoising, spike extraction, and novel microscopy methods (Zhang et al., 2023; Lecoq et al., 2021; Rupprecht et al., 2021; Zhang et al., 2024).

While experiment simulation holds significant promise, specialized software is required to attain the potential benefits. Multiple existing tools facilitate some degree of stimulation and recording of high-level population simulations (Antolík and Davison, 2013; Tomsett et al., 2015; Thornton et al., 2019; Gratiy et al., 2018; Dai et al., 2020; Dura-Bernal et al., 2019), but each has significant limitations. Many are oriented towards detailed, multi-compartment neuron models that can be hard to develop or costly to run for large populations, and none offer a full suite of ready-to-use light, opsin, and imaging models for optophysiology. Moreover, of the few that support flexible closed-loop control, none simulate the important feature of real-time processing latency for feedback control.

To address this need, we present the new open-source software Cleo: the Closed-Loop, Electrophysiology, and Optophysiology experiment simulation testbed. Cleo integrates closed-loop signal processing and virtual recording/stimulation devices with existing Brian simulator (Stimberg et al., 2019) network models to flexibly simulate a variety of experiments (see Fig. 1a). Cleo currently implements spike and approximate local field potential (LFP) recording, light and opsin models for one– and two-photon optogenetics, and two-photon calcium imaging, all with a modular design that allows for future addition of other modalities. Here we describe the design and features of Cleo, verify individual components, and validate the full system in end-to-end experiments. We further demonstrate the utility of Cleo in a variety of prospective experiments, including closed-loop inhibition of a traveling wave in sensory cortex, dynamic clamping of firing rate to disrupt visual cortex plasticity, and sharp wave-ripple evocation in the hippocampus.

## 2. Materials and Methods

### 2.1. Architecture and design rationale

In our design of Cleo, building an *in silico* experiment around an existing Brian spiking neural network model consists of (1) specifying the recording apparatus, (2) specifying the stimulation apparatus, and (3) configuring an I/O processor to control stimulation devices (see Fig. 1a, Fig. 2a). Cleo’s CLSimulator object integrates these components and orchestrates the experiment by injecting devices, running the Brian simulation, and communicating with an IOProcessor object at each time step. The IOProcessor receives measurements according to a user-specified sampling schedule and returns any updates to stimulator devices. Below, we describe the principles and assumptions that guided our modeling and software choices.

**Figure 2:**
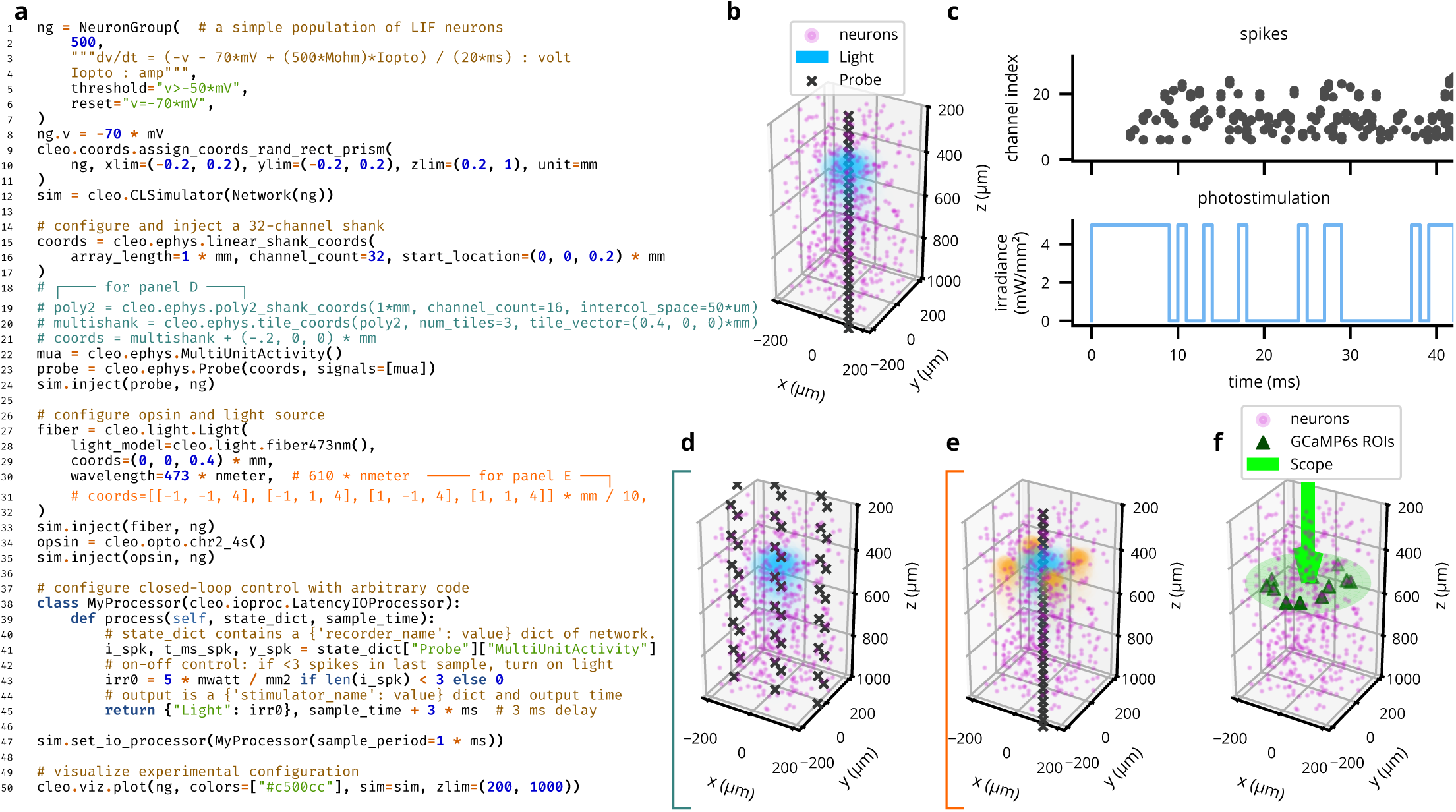
Cleo can flexibly configure electrophysiology, optogenetic stimulation, and closed-loop control with little code. (A) Code for a complete example experiment—note how few lines are needed starting with a Brian Network model. (B) Cleo provides functionality for easy visualization experiment configuration. Plot output (with slight modification) is produced by the example code in *A*. (C) Spiking data from probe and stimulation levels from fiber after running the simulation configured in *A*. (D,E) Variations in recording and stimulation devices are trivial to implement. Here, multi-shank probes and multi-site/-color stimulation are specified with coords and wavelength parameters. (F) Two-photon imaging is also trivial to configure and visualize. See Fig. 6e for a minimal code snippet.

Two factors drove our choice of recording and stimulation models to integrate into Cleo. First, because Cleo’s purpose is to simulate experiments, we focused on models at the level of accessible experimental parameters. Because parameters such as electrode location, channel count, and optic fiber depth are all defined naturally in space, Cleo’s electrode, optogenetics, and imaging modules require a spatial network model where relevant neurons are assigned *x*, *y*, and *z* coordinates. Second, we tailored Cleo to systems neuroscience models that capture mesoscale phenomena (at the circuits/population level rather than single-cell or whole-brain levels) without high degrees of biophysical realism. Specifically, Cleo was developed primarily for point neuron rather than multi-compartment, morphological neuron models. While limiting the network model space compatible with Cleo, this choice dramatically simplifies software development and reduces simulation runtime, freeing researchers to move more quickly towards the ultimate goal of informed *in vivo* experiments. This decision had consequences in our software and modeling decisions (see Sec. 2.2, Sec. 2.3, Sec. 2.4). Multi-compartment neurons are supported by Brian and could thus be better integrated into Cleo in the future. The effort required would vary from practically none in the case of spike detection without full waveforms, to moderate in extending the optogenetics and imaging APIs to allow subcellular (e.g., soma) targeting, to significant in implementing entirely new functionality such as biophysical forward modeling of extracellular potentials.

In addition to our modeling priorities, the goals of usability, flexibility, and extensibility guided our choices in software dependencies and infrastructure. Ease of use is important to make Cleo as accessible as possible, especially to researchers with primarily experimental backgrounds. This usability goal also motivated Cleo’s modular design, which allows users to add different recording or stimulation devices with little or no modification to the underlying network model, easing the burden of testing a variety of experimental configurations (see Fig. 2 for example code and visualizations). Flexibility in the underlying simulator, in addition to enabling compatibility with a wide variety of models, was a necessity for arbitrarily interacting with the simulation in a closed-loop fashion. Finally, we endeavored to make Cleo extensible so it could be adapted to use cases beyond the capabilities provided upon release, motivating the modular “plug-in” architecture that enables future incorporation of new experimental interfaces (e.g., microstimulation). In the following sections we describe the specific infrastructure and modeling choices we made in accordance with this rationale.

### 2.2. Simulator infrastructure

Other tools in the spirit of experiment simulation exist, though none with the collection of goals and functionality of Cleo. One is Mozaik (Antolík and Davison, 2013), which can manage stimulation and recording parameters as well as data and visualizations, running on the simulator backend-agnostic PyNN interface (Davison et al., 2009). It has been used to prototype and characterize advanced optogenetic control (Antolik et al., 2021; Berling et al., 2024), but PyNN does not provide an API for natively adding arbitrary differential equations to the core simulation (i.e., for features such as opsin and calcium dynamics). Three more (BioNet (Gratiy et al., 2018; Dai et al., 2020), NetPyNE (Dura-Bernal et al., 2019), and LFPy (Lindén et al., 2014; Hagen et al., 2018)) include some of the features we needed, but as front-ends to the NEURON simulator (Hines and Carnevale, 1997) they are oriented towards biophysically detailed, expensive-to-simulate neuron models. The same can be said of VERTEX (Tomsett et al., 2015; Thornton et al., 2019), which is a tool for use in MATLAB. NAOMi (Song et al., 2021) produces highly realistic two-photon calcium imaging data, but is not designed to capture other important facets of experiment simulation. See Table 1 for details.

**Table 1:** Feature comparison of experiment simulation software. †: Mozaik can record spikes from a subset of neurons selected by proximity to electrodes, but does not simulate LFP or noisy, distance-dependent spike detection with collisions.

| Software | Simulator | Ideal for point neurons | Closed-loop stimulation | Opsin kinetics models | Reconfigurable illumination | Extracellular recording | 2P imaging |
| --- | --- | --- | --- | --- | --- | --- | --- |
| Mozaik (Antolik and Davison, 2013) | Multiple (through PyNN) | ✓ | ✓ | ✓ | 1P | † |  |
| LFPy 2.0 (Hagen et al., 2018) | NEURON |  |  | ✓ |  | ✓ |  |
| BioNet (Gratiy et al., 2018; Dai et al., 2020) | NEURON |  | ✓ | ✓ |  | ✓ |  |
| NetPyNE (Dura-Bernal et al., 2019) | NEURON |  |  | ✓ |  | ✓ |  |
| VERTEX 2.0 (Tomsett et al., 2015; Thornton et al., 2019) | Custom MATLAB |  | ✓ -<br>(inflexible) |  |  | ✓ |  |
| NAOMi (Song et al., 2021) | Custom MATLAB |  |  |  | 2P |  | ✓ |
| Cleo | Brian 2 | ✓ | ✓ +<br>(w/ latency) | ✓ | 1P & 2P | ✓ -<br>(approx.) | ✓ |

Between the two most widely used spiking neural network simulators optimized for point neurons, Brian 2 (RRID:SCR_002998) (Stimberg et al., 2019) and NEST (Gewaltig and Diesmann, 2007), we chose Brian, following the example of other notable open-source projects (Evans et al., 2016; Pagkalos et al., 2023). The main advantage of Brian for our use case is the accessibility of two key features. First, custom equations (which we use to model light propagation, opsins, and calcium indicators) are used directly, whereas in NEST one must explicitly generate code using NESTML (Plotnikov et al., 2016) and install the resulting module. Second, Brian supports the execution of custom code or even arbitrary Python code directly, while this could be achieved by repeatedly stopping, modifying, and restarting the simulation or through the MUSIC multi-simulation coordination interface (Djurfeldt et al., 2010) in NEST. Nevertheless, NEST remains a powerful simulator with a large user base and many features that are not yet implemented in Brian, such as a graphical interface for network building (Spreizer et al., 2021). As such, supporting multiple simulators would greatly expand the reach of a multi-experiment simulator, but the complexity of manipulating the internals of each one led us to limit the scope of Cleo to Brian for now (see Sec. 4).

### 2.3. Optogenetics models

Cleo simulates optogenetic stimulation by combining a model of light propagation with an opsin model relating light to current. The light model is based on Kubelka-Munk light propagation, operating on the assumption that the medium is optically homogeneous and that particles are larger than the light wavelength (Foutz et al., 2012; Vo-Dinh, 2003). Cleo includes absorbance, scattering, and refraction parameters for 473-nm (blue) light as given by Foutz et al. (2012), but these are easily updated by the user for other wavelengths.

Independent of the light propagation model, Cleo provides two different opsin models. One is a four-state Markov model as presented by Evans et al. (2016). This model captures rise, peak, plateau, and fall dynamics of the photocurrent as opsins are activated and deactivated through a Markov process. By defining conductance rather than current directly, this model is also able to reproduce the photocurrent’s dependence on the membrane potential. While the four-state model fits experimental data fairly well, the code is structured so that three– or six-state models could also be easily implemented. Cleo provides parameters for channelrhodopsin-2 (ChR2) (Nagel et al., 2003), ChR2(H134R) (Nagel et al., 2005), Chrimson (Klapoetke et al., 2014), Vf-Chrimson (Mager et al., 2018), GtACR2 (Govorunova et al., 2015), and eNpHR3.0 (Gradinaru et al., 2010), as given by Evans et al. (2016) and Bansal et al. (2020b). (Bansal et al., 2020b). Users wanting to take advantage of additional optogenetic innovations such as improved channel rhodopsins (Gunaydin et al., 2010; Lin et al., 2013; Hochbaum et al., 2014; Sridharan et al., 2022; Kishi et al., 2022), chloride pumps (Chuong et al., 2014; Berndt et al., 2016) and channels (Govorunova et al., 2017), and others (Berndt et al., 2016; Vierock et al., 2021) will need to provide opsin model parameters, many of which are available in published literature (Saran et al., 2018; Bansal et al., 2020b,a; Gupta et al., 2019; Bansal et al., 2021).

However, because the Markov model depends on somewhat realistic membrane potential and resistance values, it is not well suited for many pre-existing models that do not. For example, many commonly used leaky integrate-and-fire (LIF) neurons define the membrane potential as ranging from 0 to 1, rather than –70 mV to –50 mV, rendering both the units and values (relative to the opsin’s reversal potential) incompatible. While one could adapt neuron models for compatibility with this Markov opsin model, to minimize user burden we also developed an alternative model that delivers photocurrent proportional to the light intensity at each neuron. Specifically, we offer an optional model of the opsin current described with

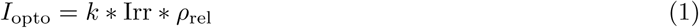

where *k* is an arbitrary gain term, Irr is the irradiance of the light source at the location of the neuron with unit mW*/*mm^2^, and *ρ*_rel_ ≥ 0 is the relative opsin expression level (the default value of 1 corresponding to the standard model fit). Note that *k* is expressed in [unit of *I*_opto_] ∗ mm^2^*/*mW, adapting to the units of *I*_opto_. This model allows users to retain their neuron model with parameters and units unchanged, since they can adapt the *k* term to whatever scale and units are needed. Preliminary experiments show that this simplified opsin model (see Fig. S3) can produce responses that are similar in many respects to those of the four-state Markov model.

In addition to options for opsin modeling, Cleo allows the user to specify both the probability that cells of a target population express an opsin and the per-cell expression level (via the afore-mentioned *ρ*_rel_ parameter). Users can thus study the impact of heterogeneous opsin expression on the outcome of an experiment. We note that this model does not describe long-term decay in opsin efficacy with prolonged stimulation.

#### 2.3.1. Multi-wavelength sensitivity

More sophisticated experimental manipulations may require the use of multiple opsins simultaneously. However, overlapping wavelength sensitivities can lead to crosstalk; i.e., a given opsin pair may not be independently controllable when light at one wavelength activates both opsins. Cleo simulates this important phenomenon using the action spectrum of each opsin. We extracted action spectra from literature (Nagel et al., 2003; Gradinaru et al., 2010; Govorunova et al., 2015; Mager et al., 2018) and represented the normalized response for stimulation of given irradiance with the factor *ε*(*λ*_other_) (Bansal et al., 2020b). For an opsin receiving light from two wavelengths, *λ*_peak_ and *λ*_other_, we then compute the effective irradiance for a given neuron as

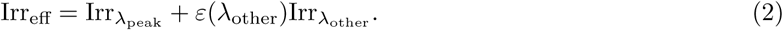

Combining irradiance linearly between light source makes the simplifying assumption that an opsin’s response to photostimulation is a linear function of irradiance (see Sec. 7.1 for details). For an example simulation of multi-wavelength, multi-opsin stimulation, see Fig. S4.

### 2.4. Electrode recording models

Because we have prioritized point neuron simulations, the electrode functionality currently implemented in Cleo does not rely on biophysical forward modeling of extracellular potentials that could only be computed from multi-compartment neurons (Pettersen et al., 2012; Buzsáki et al., 2012). Here we describe how Cleo models spiking and local field potential (LFP) recording for simpler models requiring only spikes.

#### 2.4.1. Spiking

To approximate spike recording in a lightweight manner compatible with point neuron models, Cleo implements a custom probabilistic spike detection method that does not rely on full extracellular action potential (EAP) waveforms. Our goal was to capture key features of real experiments (i.e., decaying amplitude with distance, noise, and inter-spike collisions) while keeping the simulation efficient (vectorizable), causal, simple, robust, flexible, and as realistic as possible while meeting the above criteria The core functionality of our method consists in measuring the amplitude of each spike on each channel, which decays with 1*/r*^2^ by default, where *r* is the distance from the neuron to the electrode contact (Pettersen and Einevoll, 2008) (see Fig. 3a). Measurements vary stochastically due to both background (electrical and biological) noise and intrinsic variability in spike amplitude. The user can define recording quality by setting *r*_noise floor_, the distance at which the signal-to-noise ratio (SNR) is 1; *r*_noise floor_ = 80 µm by default (Cohen and Miles, 2000). Measurements (both spikes and pure noise) that exceed a threshold *ϑ* (default 4*σ_b_*, where *σ_b_* is the standard deviation of background noise) are considered candidate threshold crossings. These threshold crossings are then sampled for “collisions”, when overlap with closely timed spikes on the same channel prevents detection (see Fig. 3b). Cleo then processes remaining spikes into multi-unit activity (MUA) output, reporting spikes detected per channel, or sorted output, where spikes detected on any channel are reported along with an index uniquely identifying the source unit (among neurons above an SNR threshold, by default 6*σ_b_*). See Fig. 3c,d for example output and Sec. 7.3 for further implementation details.

**Figure 3:**
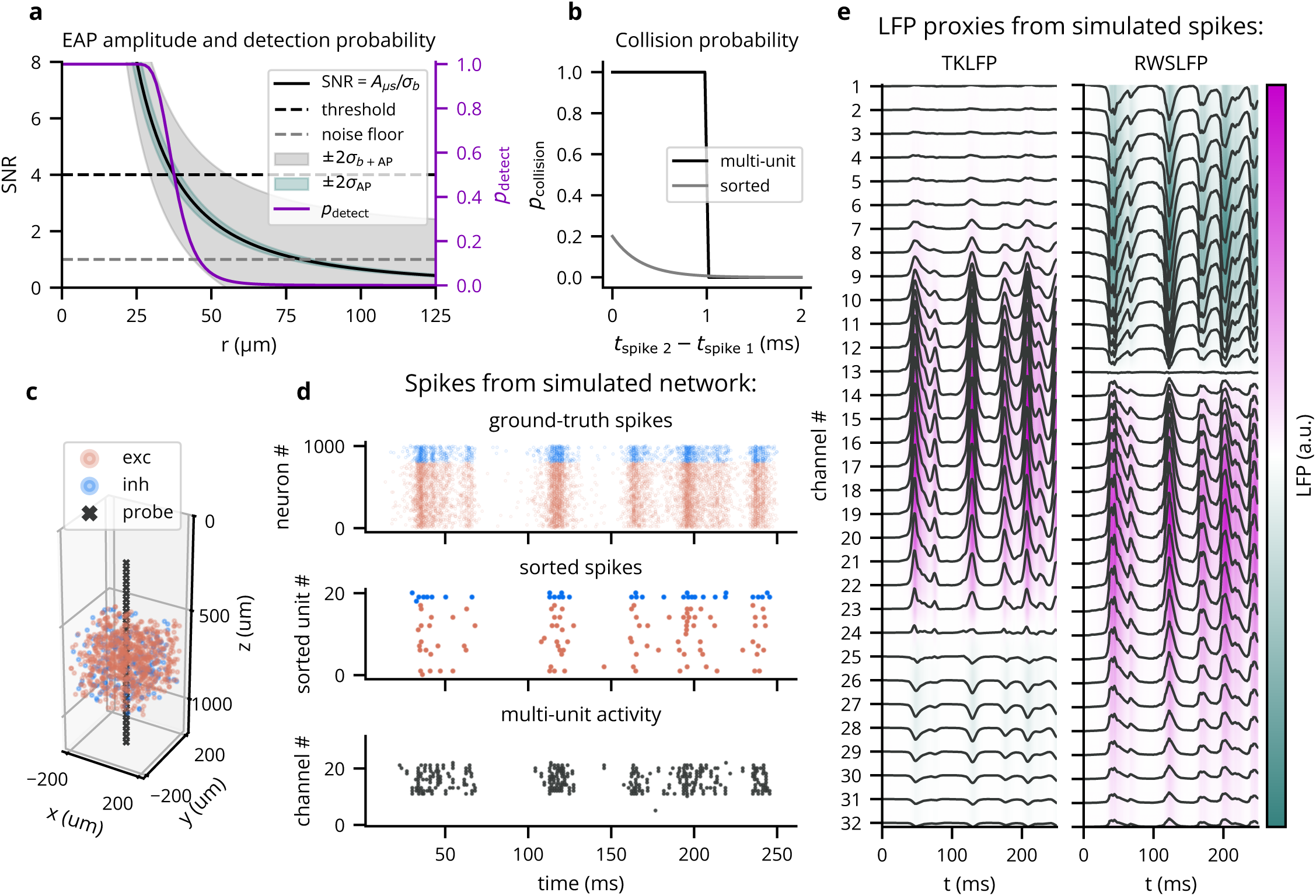
Cleo can simulate spike and LFP recording from point neuron network models. (A) Default parameters/functions of the threshold crossing stage of the probabilistic spike detection model. SNR describes EAP amplitude as a function of distance from the neuron to the electrode *r*. Shading describes ±2*σ* variability in the Gaussian-distribution EAP amplitudes. *p*_detect_ reflects the probability a measured amplitude crosses the threshold on a single channel. (B) Default collision probability functions for the collision sampling stage of the probabilistic spike detection model. (C) A plot generated by Cleo showing the positions of neurons and electrode contacts in a sample excitatory/inhibitory network. The contacts emulate a 32-channel linear NeuroNexus array. (D) Spiking activity recorded in the setup shown in *C*. Ground-truth spikes are taken directly from the Brian simulation, while sorted and multi-unit spikes are recorded with Cleo using default parameters. (E) The two LFP proxy signals provided by Cleo, recorded from the same simulated network/activity in *C* /*D*.

**Figure 4:**
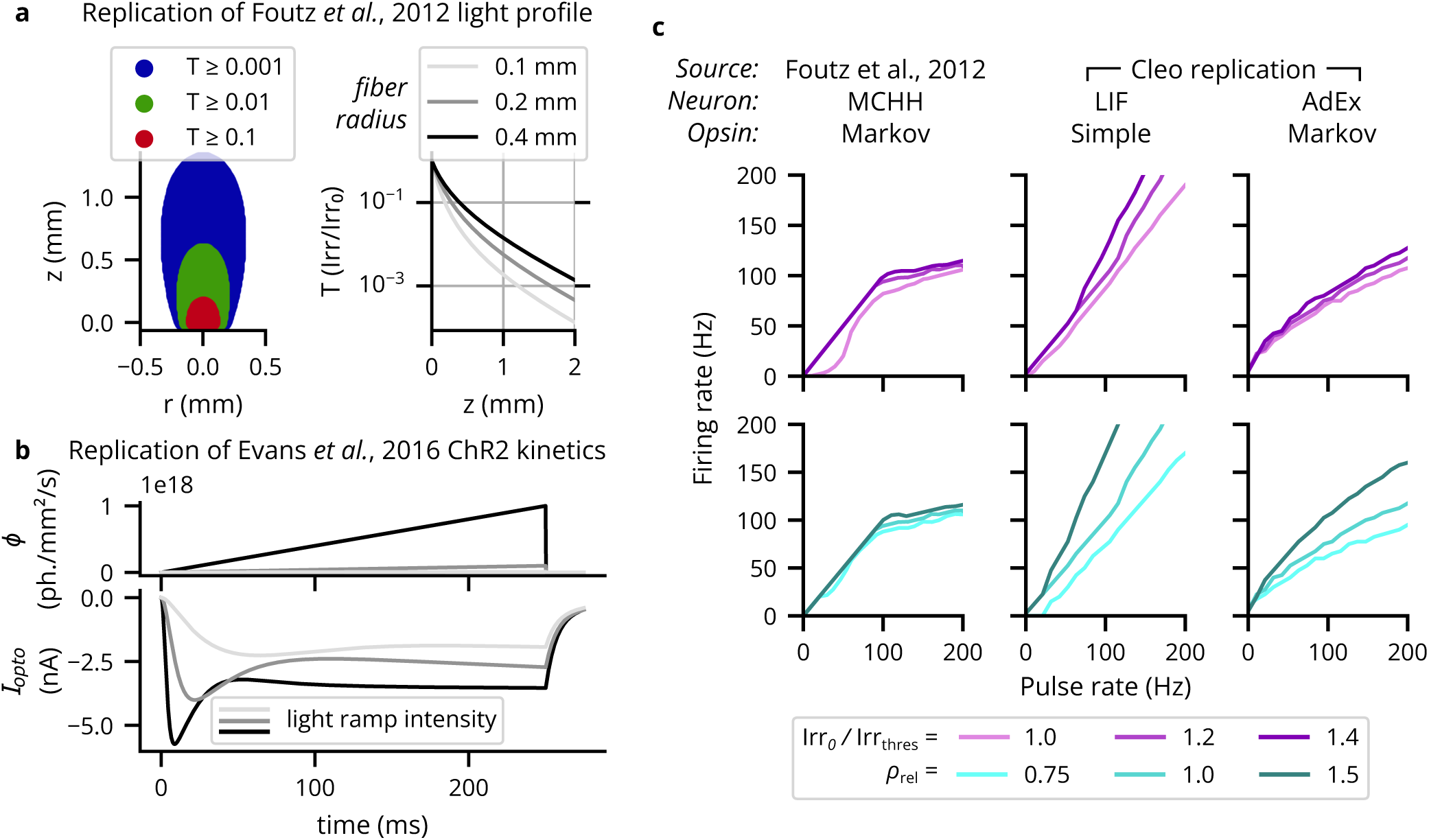
Verification of the optogenetics module. (A) Cleo exactly replicates the theoretical light profile from (Foutz et al., 2012). Left: Light transmittance *T* as a function of radius and axial distance from the optic fiber tip (cf. Figure 2a from (Foutz et al., 2012)). See Fig. S1a for more detail. Right: Light transmittance *T* as a function of distance *z* straight out from the fiber tip for different optic fiber sizes (cf. Figure 2b from (Foutz et al., 2012)). (B) Photocurrent *I*opto for ramping light of different intensities (cf. Fig. S2). (C) Different neuron/opsin model combinations replicate detailed simulations with varying degrees of success. We plot neuron firing rates in response to optical stimulation with 5-ms pulse frequencies ranging from 1 to 200 Hz. The left column re-plots data from (Foutz et al., 2012), using the same Markov opsin model implemented by Cleo with a multi-comparment Hodgkin-Huxley neuron model. The middle column shows results for an LIF neuron with a simple opsin, and the right column for an AdEx neuron with a Markov opsin model. The top row shows results for different light intensities: 100%, 120%, and 140% of the threshold for producing a single spike with a 5-ms pulse. The bottom row shows results for different expression levels relative to the default, *ρ*_rel_. See Fig. S1b for more neuron model-opsin combinations.

#### 2.4.2. LFP

To approximate cortical LFP without resorting to morphological neurons and biophysical forward modeling, we implemented two LFP proxy signals that can be computed from point neuron simulations (see Fig. 3e). The first approximates the per-spike contribution to LFP with a delayed Gaussian kernel, where amplitude and delay depend on the position of the neuron relative to the electrode, as well as cell type (excitatory or inhibitory) (Telenczuk et al., 2020). We hereafter refer to this proxy signal as Teleńczuk kernel LFP (TKLFP). Default parameters (taken from the original study) were estimated from human temporal cortex experimental data and from hippocampus simulations. As the authors indicate, parameters may need refinement on a per-region basis. While the original study included reference peak amplitude (*A*_0_) values at just four cortical depths, we inferred these values for arbitrary depths by performing cubic interpolation on reported data (see Figure 5 in (Telenczuk et al., 2020)) and assumed that this profile dropped to zero at 600 µm below the soma and 1000 µm above.

**Figure 5:**
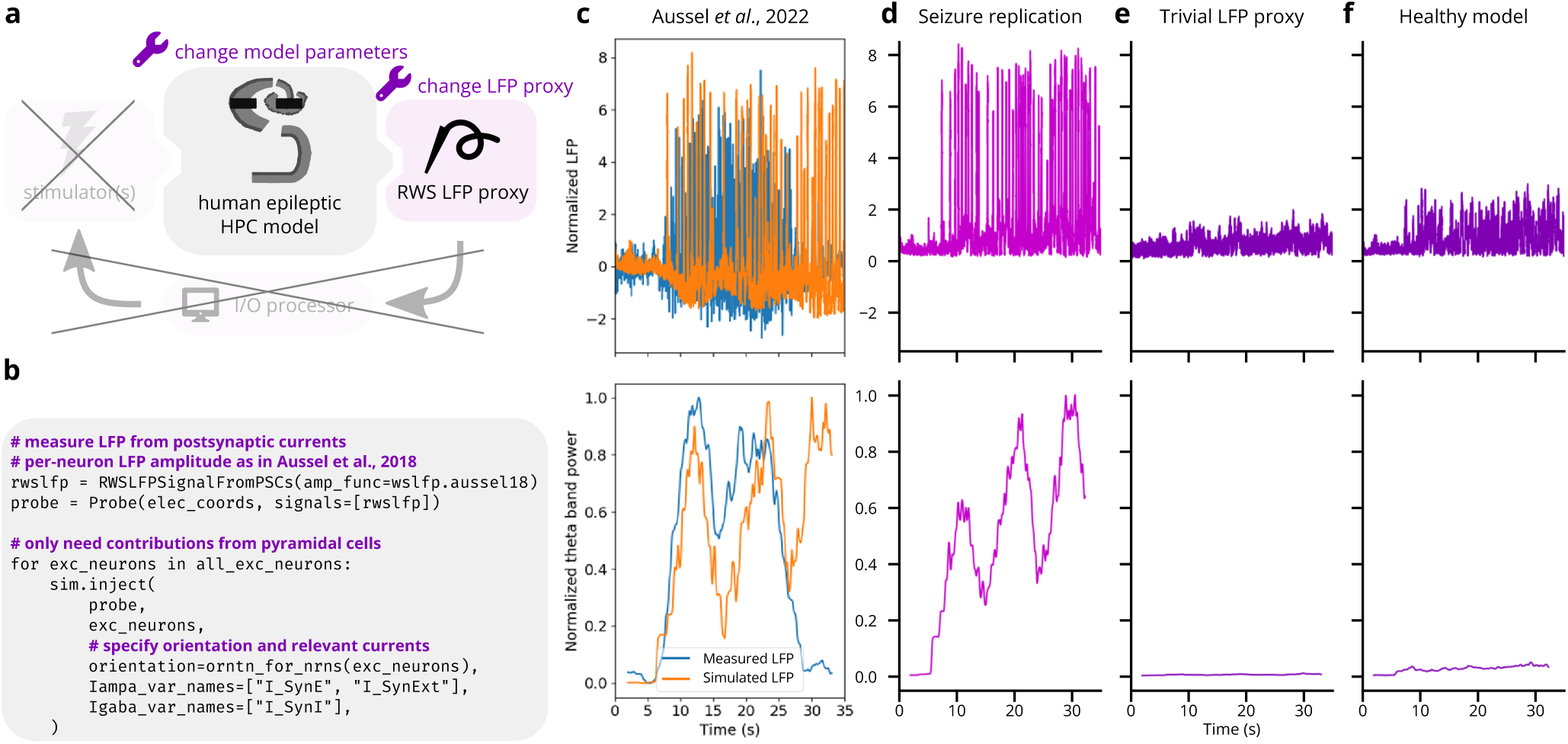
Reproduction of electrophysiological recordings of epileptiform hippocampus activity (Aussel et al., 2022). (A) Schematic of experiment setup. LFP is recorded from a hippocampal model (Aussel et al., 2022), and ablations of both the model parameters and LFP output serve as negative controls. (B) Minimal code required to record LFP from the existing model. This replaces hundreds of lines in the original model code. (C) Top: Experimental and simulated LFP (estimated from summed synaptic currents). LFP is normalized to have a peak of 1 during the first 5 seconds of the simulation. Bottom: theta band power, normalized by the peak value. Theta band power was computed using SciPy’s spectrogram function with a Tukey window of width 3.906 sec, *α* = 0.25, and overlap width of 3.809 sec. Image used under the CC BY 4.0 license. (D) Replication of *C* via Cleo’s RWSLFP recording. Theta power is normalized by the peak value. (E) Same as *D*, but with the average model input serving as an ablated LFP output. Theta power is normalized by the peak in *D*. (F) Same as *D*, but with model parameters corresponding to a healthy, rather than an epileptic state. Theta power is again normalized by the peak in *D*.

Cleo also provides the Reference Weighted Sum of postsynaptic currents LFP proxy (RWSLFP) (Mazzoni et al., 2015), which fits the forward model LFP well (*R*^2^ *>* 0.9) for standard pyramidal cell morphologies when network activity and recording location yield a sufficiently large signal. This method sums AMPA and GABA currents onto pyramidal cells, each current with a different weight and time delay. The amplitude of the signal is then determined by the axial and lateral recording distances, relative to pyramidal cells’ apical dendrites. To support arbitrary recording locations, we interpolated and extrapolated this amplitude profile as given in Figure 2B of the original publication. We did this by fitting a scaled beta distribution kernel at each radial distance and interpolating linearly between these fits. Because these signal amplitudes were evaluated by summing currents over a population distributed within a 250 µm-radius cylinder, Cleo supports arbitrary morphologies by providing an alternate amplitude profile optimally scaled such that the sum of individual neurons’ contributions is close to the population profile. We also include a scaled version of the closed-form per-neuron contribution as given by Aussel et al. (2022).

A major difference between the two methods is that TKLFP is computed from spikes alone, while RWSLFP requires synaptic currents. Continuing in the spirit of supporting simplistic network models, Cleo provides the option to synthesize synaptic currents instead of simulating their dynamics by convolving spikes with a biexponential kernel (see Eq. (5.34) in (Dayan and Abbott, 2005)), requiring only that the user specify which synapses mediate these spikes. The basis in currents allows RWSLFP to better capture high-frequency signals deriving from subthreshold activity (see Fig. S11).

As there was no publicly available code flexibly implementing these methods, we created, tested, and documented standalone implementations in the tklfp and wslfp Python packages (Johnsen, 2022; Johnsen et al., 2024). The authors’ goal of lowering the cost of LFP simulation is thus aided as their methods are easily accessible for the first time, for use inside or outside Cleo.

Besides these two methods, we note that another, hybridLFPy (Hagen et al., 2016), has a similar goal of approximating LFP from point neuron models. This method is more realistic than the two described here, using the activity of a point neuron network and propagating the resulting spikes through a mutually unconnected network of morphological neurons, which are then used to compute the LFP in the forward model. Incorporating this method into Cleo would require instantiation of and bidirectional communication with a twin network of morphological neurons, which we deemed too costly and complex to align with our principal goal of making Cleo relatively lightweight. This should be possible, however, and could be a valuable addition to Cleo in the future. Short of this, the validity of the TKLFP and RWSLFP methods can be assessed from the reports describing them initially (Telenczuk et al., 2020; Mazzoni et al., 2015). Such an approach would be the best way to evaluate the accuracy of the TKLFP and RWSLFP modules; such an evaluation is outside the scope of this work and we refer readers to their source publications to judge their validity.

### 2.5. All-optical control

#### 2.5.1. Two-photon microscopy

Cleo simulates microscopy by taking microscope location, image width, focus depth, and soma size, and selecting neurons with a cross section in the plane of imaging. Calcium traces are generated for the given regions of interest (ROIs), adding Gaussian noise of standard deviation *σ*_noise_ that depends both on the indicator and on the size of the soma cross section in focus. We model noise as Gaussian as a consequence of the central limit theorem, since the ROI measurement is a sum of per-pixel stochastic measurements (Lütcke et al., 2013) (see Fig. S7). Accordingly, we scale *σ*_noise_ with 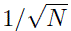, where *N* is the number of visible pixels relative to the maximum (when the center of the soma lies exactly on the focal plane; see Fig. 6b,c, Fig. S8). Signal strength is proportional to expression, denoted as *ρ*_rel_ as with opsins (see Fig. 6c). Thus, for ROI *i*:

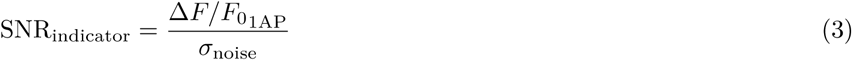

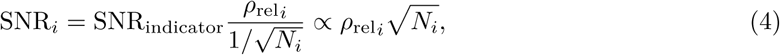

where Δ*F/F*_01AP_ is the Δ*F/F*_0_ peak after a single action potential. Δ*F/F*_01AP_ and *σ*_noise_ are indicator-specific and taken from Dana et al. (2019) and Zhang et al. (2023). ROIs with signal-to-noise ratio (SNR) above a specified cutoff are selected for recording. Each parameter can be set independently for each injection. Expression levels can additionally be set per neuron; per-neuron soma size variation is not currently supported but could be added in the future.

**Figure 6:**
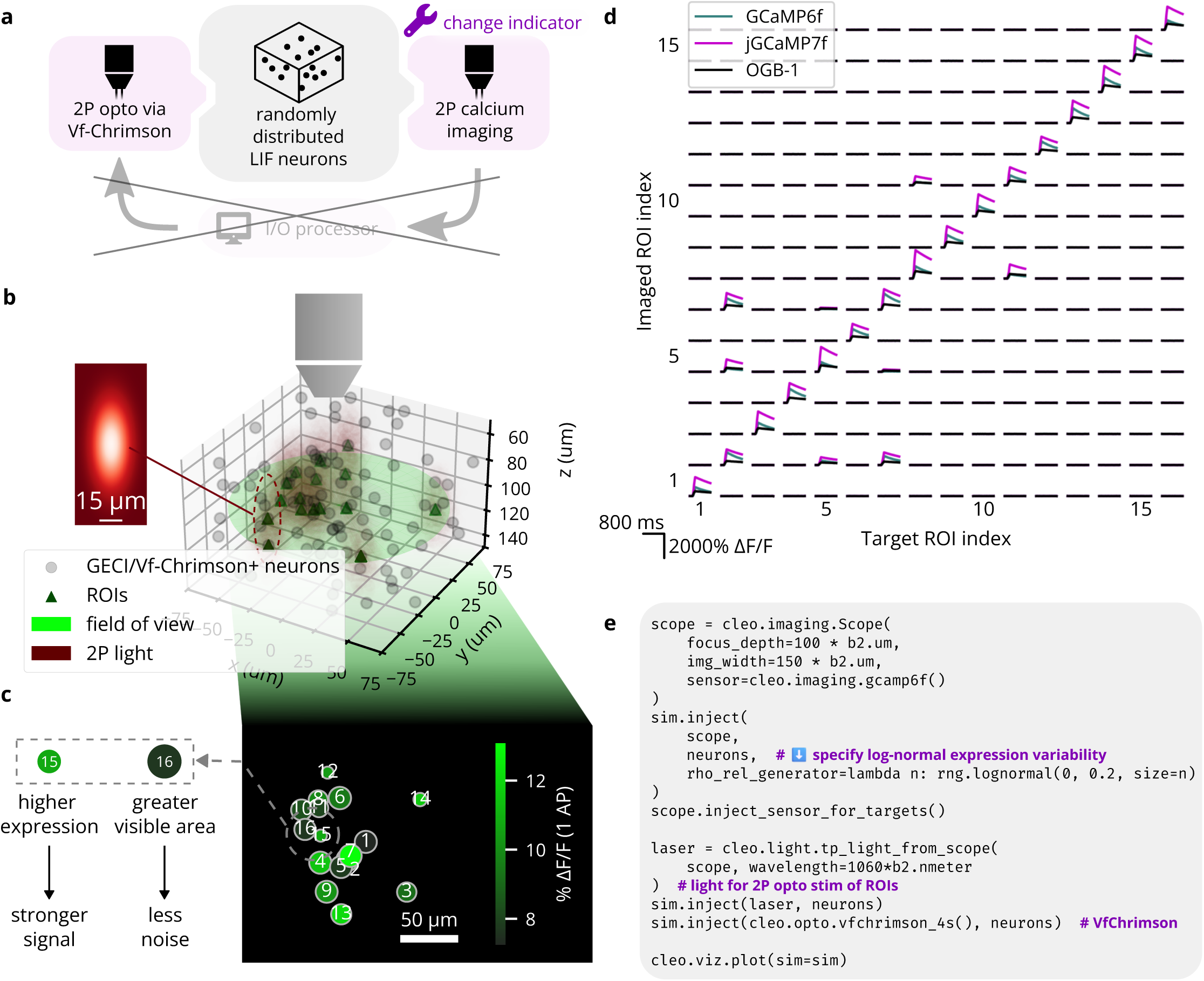
Reproduction of an end-to-end all-optical control experiment, after Figure 3 of Rickgauer et al. (2014). (A) A schematic of the experiment configuration. Different calcium indicators are simulated to demonstrate Cleo’s capability to aide experimental design. (B) A 3D plot of the model spiking neural network with the microscope’s field of view visualized. Dark red ellipsoids depict laser light intensity around targeted neurons. *Inset*: A heatmap visualization of the Gaussian point spread function defining light intensity around each 2P stimulation target; cf. Figure 3b of Rickgauer et al. (2014). The *x* and *y* axes correspond to lateral and axial axes, respectively. (C) 2D image as seen by the microscope; cf. Figure 3c of Rickgauer et al. (2014). Size represents how much of each ROI is visible, i.e., how well centered it is on the focal plane. Color indicates signal strength, as determined by expression levels. (D) Results from the simulated all-optical experiment; cf. Figure 3c of Rickgauer et al. (2014). Microscopy and photostimulation are configured as in *B*, performing calcium imaging using a model of the OGB-1, GCaMP6f, and jGCaMP7 indicators (Kerr et al., 2005; Chen et al., 2013; Dana et al., 2019). Each ROI is targeted one at a time (represented in each column), receiving 10 pulses of 2 ms width at 100 Hz. The recorded calcium trace of each ROI is shown in each row. Off-target effects can be seen for neurons that are close together (e.g., 8 and 11). (E) Minimal code example for configuring all-optical control, including the microscope, opsin, and calcium indicator. rho_rel refers to the expression level.

#### 2.5.2. Calcium indicator model

Cleo simulates intracellular calcium concentration dynamics using a biophysical model described previously in literature (Song et al., 2021; Helmchen and Tank, 2015; Lütcke et al., 2013):

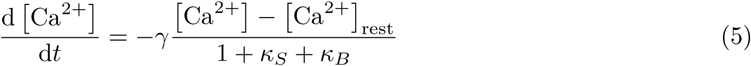

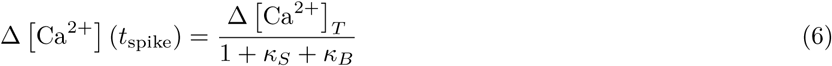

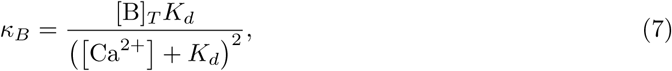

where *γ* is the clearance rate, *κ_S_* is the endogenous Ca^2+^ binding ratio, *κ_B_* is the Ca^2+^ binding ratio of the exogenous buffer (the indicator), *K_d_* is the indicator dissociation constant, [B]*_T_* is the total intracellular indicator concentration, and Δ [Ca^2+^] is total [Ca^2+^] increase per spike. Following Song et al. (Song et al., 2021), Ca^2+^ (*t*) is then convolved with a double exponential curve *h*(*t*) to obtain [CaB_active_], reflecting the response kinetics (such as binding and activation) not accounted for by binding affinity *K_d_* alone (Badura et al., 2014):

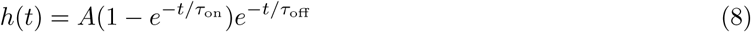

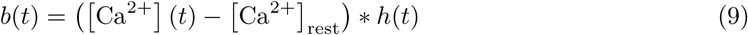

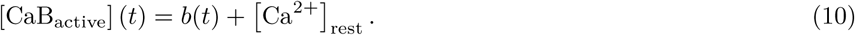

Parameters *A, τ*_on_*, τ*_off_ are indicator-specific. This convolution is approximated as integration of an ODE for ease of simulation (see Sec. 7.2).

Δ*F/F*_0_ is then computed from [CaB_active_] using a Hill equation nonlinearity and subtracting the baseline value to produce Δ*F/F*_0_ = 0 when [CaB_active_] = [Ca^2+^]_rest_:

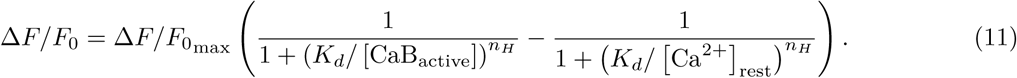

Parameter values for various genetically encoded calcium indicators (GECIs) are taken from the NAOMi simulator (Song et al., 2021). For an example of simulated traces, see Fig. S5.

#### 2.5.3. Two-photon photostimulation

Cleo simulates two-photon (2P) photostimulation using the same opsin models previously described (Sec. 2.3) by modeling focused laser illumination. As is commonly reported in 2P experiments, laser power is used to define stimulation intensity. We convert from power to irradiance (needed for opsin models) by dividing by soma area (Ronzitti et al., 2017), assuming a diameter of 20 µm. We then model off-target effects using a Gaussian ellipsoid point spread function with *σ*_axial_ *> σ*_lateral_, as reported in literature (Prakash et al., 2012; Rickgauer et al., 2014; Packer et al., 2015; Chen et al., 2019) (see Fig. 6b). When targeting cells identified by the microscope, the laser is focused on the plane of imaging, such that the farther off-plane cells, the weaker they are stimulated. Morphological factors of 2P photostimulation such as membrane-boundedness of the opsin and differential expression between the soma and processes are not modeled.

### 2.6. Latency model

To simulate the effects of real-time compute latency, Cleo provides a LatencyIOProcessor class capable of delivering control signals after arbitrary delays. It does this by storing the outputs calculated for every sample in a buffer along with the time they can be delivered to the network. For example, if a sample is taken at time *t* = 20 ms and the user wishes to simulate a 3 ms delay, the control signal and output time (23 ms) are stored in a buffer which the simulation checks at every time step. As soon as the simulation clock reaches 23 ms, the control signal is taken from the buffer and applied to update the stimulator devices. Because the user has arbitrary control over this latency, they can easily simulate the effects of closed-loop experimental constraints. For example, one could use probabilistic delays to assess the effect closed-loop algorithms with variable round-trip times between measurement and stimulation. By default, LatencyIOProcessor samples on a fixed schedule and simulates processing samples in parallel (i.e., the computation time for one sample does not affect that of others). This and the other sampling schemes Cleo provides are illustrated in Fig. S9.

### 2.7. Neo export

To maximize compatibility with existing data analysis packages and pipelines, Cleo supports data export using Neo (RRID:SCR_000634), a Python package providing an in-memory representation of neuroscience data and the ability to read dozens of file formats (Garcia et al., 2014; Davison et al., 2022). Analysis code developed for experiments could thus be reused for simulated data, and vice versa.

### 2.8. Computing environment and performance

Experiments (described in Sections 3.2 and 3.3) were run in one of three environments. The first is the Georgia Tech Partnership for Advanced Computing Environment (PACE) Phoenix cluster with 64 GB RAM, dual Intel Xeon Gold 6226 CPUs @ 2.7 GHz (24 cores/node), DDR4-2933 MHz DRAM, and Infiniband 100HDR interconnect. The second is a Dell consumer laptop with an Intel i9-9980HK CPU @ 2.40 GHz (8 cores) and 32 GB RAM. The third is a Puget Systems desktop computer with Intel Xeon W-2255 @ 3.7GHz 10-core CPU and 128 GB RAM. Note that Cleo simulations are not parallelized because the arbitrary Python code implementing many features precludes compilation. All experiments were run with Cleo version 0.18.1. See Table 2 for a list of the computing environment, model details, and runtime of each experiment.

**Table 2:** Experiment computation details. Runtimes describe individual conditions/trials, rather than the entire experiment. VE: validation experiment, PE: prospective experiment, LIF: leaky integrate-and-fire, HH: Hodgkin-Huxley. †: For the epileptic model; the healthy model is as PE3.

| Experiment | Computer | No. neurons | Neuron model | No. populations | No. synapses | Sim. time | Approx. runtime |
| --- | --- | --- | --- | --- | --- | --- | --- |
| VE1: HPC seizure recording | Desktop | 19,820<sup>†</sup> | HH | 8 | $\sim 1.1 \times 10^7$<sup>†</sup> | 35 s | 160 min |
| VE2: All-optical control | Dell laptop | 100 | LIF | 1 | 0 | 800 ms | 15/1.5 s with/without imaging |
| VE3: Bidirectional optoclamp | Dell laptop | 1,000 | LIF | 2 | $\sim 1 \times 10^5$ | 90 s | 7 min |
| PE1: Traveling wave rejection | Dell laptop | 12,500 | LIF | 2 | $\sim 7.4 \times 10^4$ | 15 ms | 10 s, including setup |
| PE2: V1 plasticity disruption | PACE | 694 | LIF | 5 | $\sim 4.0 \times 10^5$ | 137 s | 53/38 min with/without Cleo |
| PE3: SWR evocation | Desktop | 33,200 | HH | 8 | $\sim 2.2 \times 10^7$ | 400 ms | 3 min with/without opto |

### 2.9. Software and code accessibility

Cleo source code is hosted on GitHub at https://github.com/siplab-gt/cleo. Documentation, includ-ing an overview, tutorials, and API reference, can be found at https://cleosim.readthedocs.io. Code for experiments can be found at https://github.com/siplab-gt/cleo/tree/master/notebooks, https://github.com/siplab-gt/cleo-traveling-wave-rejection, https://github.com/siplab-gt/cleo-v1-plasticity-expt, and https://github.com/siplab-gt/cleo-hpc-experiments.

### 2.10. Feedback control

Validation experiment 1 used proportional-integral (PI) control and firing rate estimation as described in (Newman et al., 2015). Prospective experiment 2 used PI control and exponential firing rate estimation as described in (Bolus et al., 2018). Cleo provides implementations of these, which can be found in the cleo.ioproc module. Prospective experiment 3 used a standard linear quadratic regulator (LQR) approach as described in (Bolus et al., 2021) and implemented in the ldsCtrlEst C++ library v0.8.1 (Bolus et al., 2022). ldsCtrlEst is part of CLOCTools (Willats et al., 2022; Johnsen, 2022), a larger collection of algorithms and utilities for implementing closed-loop optogenetics in real-time lab experiments. Prospective experiment 3 also used a custom implementation of model-predictive control (MPC). We added 3 and 6 ms of latency to LQR and MPC, respectively, to simulate computation time. For details on model fitting and control parameters, see Sec. 7.5.

## 3. Results

We demonstrate the utility of Cleo with a variety of different results. First, we validate output from the optogenetics and LFP recording modules by comparing to data from published literature. This confirms that these nontrivial models are suitable for integration into larger simulations. Next, to establish the validity of combining multiple models into the unified simulation of a complete experiment, we compare the results of three end-to-end validation experiments to published data for various experimental paradigms. Finally, we provide examples of how Cleo can be used to prototype novel closed-loop optogenetic techniques in three prospective experiments using previously published network models. Table 2 describes the runtime of each experiment.

### 3.1. Component verification

#### 3.1.1. Optogenetics model verification

To verify Cleo’s light and opsin model implementations, we first reproduced a previously reported optic fiber light transmission model (Foutz et al., 2012). The model defines transmittance *T* as the proportion of irradiance at a given point Irr to the irradiance at the fiber tip Irr_0_. Fig. 4 demonstrates that Cleo’s transmittance model corresponds to previously reported results as a function of radius and axial distance from the optic fiber tip (cf. our panel *A* and Figure 2 of (Foutz et al., 2012)). See also Fig. S1a. Validating the four-state opsin kinetics model, we also reproduced the ChR2 photocurrents in response to ramping light stimuli of varying intensities (Evans et al., 2016) (cf. our panel *B* and Fig. S2). These results all match the source publications exactly, as they are all evaluations/simulations of the same equations.

To test how well simplified models produce realistic firing patterns on long timescales, we also compared pulse rate to firing rate for a variety of light intensities and opsin expression levels as simulated with multi-compartment Hodgkin-Huxley neurons by Foutz et al. (2012). We used combinations of leaky integrate– and-fire (LIF) and adaptive exponential integrate-and-fire (AdEx) (Fourcaud-Trocmé et al., 2003) neuron models, along with proportional current and Markov opsin models. Parameters for each model combination were obtained via Bayesian optimization, minimizing the sum of the squared error across conditions between the obtained firing rates and the target firing rates from the detailed simulations (see Sec. 7.4 for details). As expected, the different model combinations behave differently and none reproduce exactly more detailed biophysically realistic simulations (see Fig. 4c, Fig. S1b). While they all reproduce a linear relationship at low pulse rates, the more realistic model AdEx/Markov model combination is able to capture the sublinear relationship at higher pulse rates (though without a sharp bend), while the LIF/simple combination fails to do so. This both demonstrates the potential of point neuron model to substitute morphological neuron models and highlights the importance of selecting models that are detailed enough to capture phenomena of interest.

#### 3.1.2. LFP model verification

In addition to providing unit tests in the Cleo, wslfp, and tklfp codebases, we verified Cleo’s LFP output by comparing to previously published results. To test the Teleńczuk kernel LFP approximation module, we reproduced the demo presented by (Telenczuk et al., 2020) and found that Cleo’s output was essentially identical (see Fig. S10a). Here we expect a close match using the same equations and source data, but not identical because we corrected a few minor implementation errors. We also compared TKLFP and RWSLFP output of the hippocampus model to its summed synaptic current LFP proxy and found them all to be qualitatively similar (see Sec. 3.3.3, Fig. S10b) (Aussel et al., 2018). We do not expect a quantitative match in this case because each proxy signal is derived from different equations/parameters. Here and in further comparisons (see Fig. S11), we confirmed that TKLFP underrepresents high-frequency components compared to RWSLFP, as reported in the original publication. We also find its sign inverts at a depth other than that predicted by detailed biophysical modeling, namely, around the midpoint of pyramidal cell dipoles (Mazzoni et al., 2015). These evaluations suggest that the methods are implemented correctly and can thus be applied to a variety of modeling applications, subject to the limitations described by their authors.

### 3.2. End-to-end validation experiments

#### 3.2.1. Validation experiment 1: LFP recording of epileptiform hippocampus activity after Aussel et al

We illustrated Cleo’s utility in simulating electrophysiology experiments by replicating epileptiform activity recorded from the human hippocampus (see Fig. 5a). We used the model first described by Aussel et al. (2018) and adapted to model epilepsy (Aussel et al., 2022). In the epileptic setting it features a network of 19,820 point neurons with Hodgkin-Huxley dynamics and ∼1.1 × 10^6^ total synaptic connections. Excitatory and inhibitory populations are modeled for each of the entorhinal cortex, dentate gyrus, CA3, and CA1 regions, with cell locations and connection probabilities derived from anatomy; the model extends roughly 6 × 10 × 15 mm. The epileptic model modifies the “wakefulness” setting of the original model by increasing monosynaptic connection probability 50%, simulating sclerosis and sprouting, raising the equilibrium potential of potassium channels, and increasing the time constant for chloride channels. Seizures are simulated by delivering stochastic spike trains derived from stereoelectroencephalography (SEEG) recordings in three regions afferent to entorhinal cortex: the prefrontal cortex, the lateral temporal lobe, and the temporal pole. Electrodes in this model are distributed in two 0.8 cm diameter cylinders of 144 contacts each, one in the entorhinal cortex and one in the CA1 region; LFP is computed by taking the difference in the mean signal of each cylinder. The authors show that when parameters are tuned to unhealthy states, the model exhibits epileptiform activity similar to the SEEG data, especially in theta band power (see Fig. 5c).

We wrapped this model with Cleo, delivered the inputs derived from afferent brain area recordings, and recorded LFP with electrodes in the same location as in the experiment. Cleo’s RWSLFP output reproduces the epileptiform activity present in the original simulation, which in turn reproduces the rhythmic amplification of theta-band activity seen in the experimental data. This suggests Cleo can usefully simulate electrophysiology experiments provided a satisfactory spiking neural network model (see Fig. 5d). We do not expect a perfect match to the source publication’s simulation because the input in our simulation was not identical—rather, each simulation sampled a spike train independently from the same processed SEEG data. Notable differences could thus be seen between simulations, although the three-peak theta power pattern remained consistent. Other differences in the LFP signal can be attributed to differences between our RWSLFP implementation and that of Aussel *et al*.—namely in the spatial amplitude profile, the time delay for AMPA current contributions *τ*_AMPA_, and weight of GABA current contributions *α*. The latter, in their case *α* = 1 while we followed Mazzoni *et al*.’s *α* = −1.65, explains why their signal goes negative while ours does not, keeping in mind that LFP(*t*) ∝ AMPA(*t* − *τ*_AMPA_) − *α* GABA(*t*). The fact that Cleo’s output does not match the original two-peak theta pattern highlights that a simulated experiment can only be as accurate as the core network model.

To show this result was nontrivial, we also ran the same simulation with a naïve LFP model (using the average SEEG input instead of RWSLFP as the proxy signal) and an altered network (using healthy rather than epileptic model parameters). These ablations (see Fig. 5e,f) failed to produce the heightened signal and theta band power seen in the original data, suggesting that the accuracy of both the network and the LFP model play an important role in replicating the experiment.

#### 3.2.2. Validation experiment 2: All-optical stimulation and recording of individual neurons after Rick-gauer et al

To validate Cleo’s simulation of two-photon, all-optical stimulation and recording, we reproduced the data presented in Figure 3 of Rickgauer et al. (2014), where individual neurons are controlled. We distributed 100 mutually unconnected LIF neurons randomly in 150 × 150 × 100 µm 3D space and configured a microscope to record at a depth halfway through the population with a 150 µm-wide image. Using Cleo’s default microscopy settings, ROIs with SNR ≥ 1 were identified. More modern molecular tools (and the older OGB-1) available in Cleo are substituted for the original GCaMP3/C1V1 indicator/opsin setup. One of three calcium indicators (OGB-1, GCaMP6f, or jGCaMP7 (Kerr et al., 2005; Chen et al., 2013; Dana et al., 2019)) and the Vf-Chrimson opsin (Mager et al., 2018) were injected and each neuron was stimulated with 10 pulses of 2 ms width at 100 Hz. 1060 nm light at 5 mW power (tuned to produce roughly one spike per pulse) was used for stimulation. The resulting calcium traces in Fig. 6d reproduce the most important qualities of Rickgauer et al. (2014), Figure 3, namely heterogeneity in signal and noise strength and independent stimulation of neurons, limited by spatial proximity. With regards to the latter, we see off-target ROIs respond significantly but more weakly than nearby targeted ROIs, as expected.

#### 3.2.3. Validation experiment 3: In vitro optoclamp after Newman et al

Demonstrating Cleo’s ability to capture salient features of closed-loop optogenetic control experiments, we reproduced the “optoclamp” experiment of Newman et al. (2015) on cultured neurons. We simulated an E/I leaky-integrate-and-fire (LIF) network (Brunel, 2000) of 800 excitatory and 200 inhibitory cells randomly distributed in a 2 mm diameter disc with no thickness. Inhibitory weights were tuned to overpower excitatory weights, creating a network-wide bursting behavior. Specifically, the inhibitory weight multiplier *g* to *g* = 1.875*g*_balanced_, where *w_e_* = *g*_balanced_ *w_i_*. The multi-electrode array (MEA) had 60 contacts distributed with 200 µm spacing as depicted by Newman et al. (2015) and was configured to produce sorted spikes in real time. ChR2(H134R) (Nagel et al., 2005) and eNpHR3.0 (Gradinaru et al., 2010) were used as the excitatory and inhibitory opsins, respectively, injected with lognormal-distributed expression levels. These were targeted with uniform 465 nm and 590 nm light, respectively (see Fig. 7a). A proportional-integral (PI) controller as described by Newman et al. (2015) determined these light levels to clamp firing rates to different target values. Detailed parameter settings can be found in the code repository.

**Figure 7:**
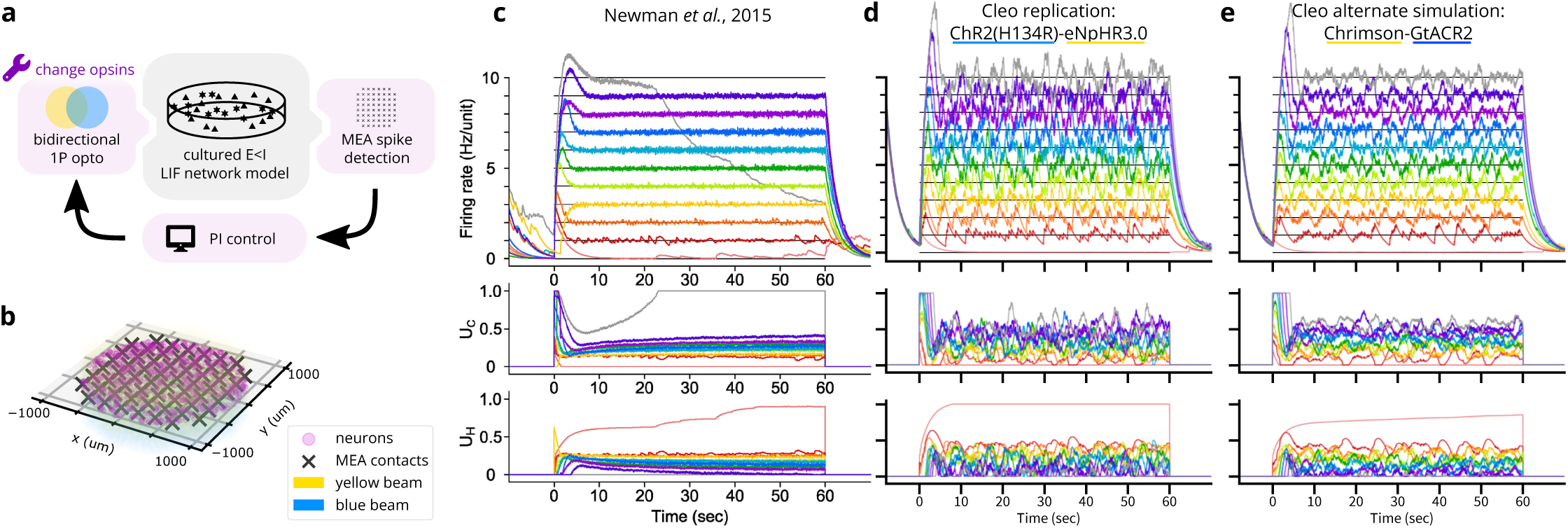
Reproduction of an end-to-end optogenetic feedback control (“optoclamp”) experiment (Newman et al., 2015). (A) Schematic of the experimental setup. Different opsin pairs are simulated to show Cleo’s capability to aide experimental design. (B) 3D plot of network model, multi-electrode array, and light configuration. (C) Experimental data from Figure 2A of Newman et al. (2015), showing firing rate (top), ChR2(H134R) control signal (*U_C_*, middle), and eNpHR3.0 control signal (*U_H_*, bottom) for each of 11 target firing rates, each marked with a different color. Image used under the CC BY 4.0 license. (D) Replication of *B* in a Cleo simulation. (E) Same as *D*, but with the Chrimson-GtACR2 opsin pair instead.

Our simulation (see Fig. 7c) reproduces key features of the experimental data (Fig. 7b) such as the initial overshoot/settling phase and the controller’s successful clamping of firing rate. Finer details such as noise levels, post-inhibition rebound, and the increase in required stimulation over time were not reproduced. These can be explained by inaccuracies in the SNN model; namely, the simple E/I LIF network lacked adaptive or homeostatic mechanisms, and a number of parameters such as synaptic weights and opsin expression levels were not finely tuned.

We also ran the experiment with an alternate opsin pair, Chrimson (Mager et al., 2018) and GtACR2 (Gov– orunova et al., 2015), which required tuning a number of parameters differently to achieve similar results. The blue light was set to 450 nm wavelength to minimize activation of Chrimson, but control gains still needed to be adjusted to prevent Chrimson from overpowering GtACR2. Pulse frequency also needed to be increased to enable Chrimson to drive firing activity fast enough for higher target rates, presumably due to its faster off-kinetics. This process provides a glimpse into the difficulties of tuning closed-loop stimulation and shows how Cleo could be used to help design robust experiments.

### 3.3. Prospective experiments

#### 3.3.1. Prospective experiment 1: Closed-loop rejection of an S1 traveling wave

We implemented a rodent primary somatosensory cortex (S1) traveling wave model (Moldakarimov et al., 2018) in Brian to demonstrate Cleo’s capabilities for simulating an event-triggered closed-loop control experiment. The rodent S1 model includes 10,000 excitatory and 2,500 inhibitory neurons distributed evenly in a 5 mm × 5 mm grid. Synapses include ∼7 × 10^4^ weak local connections and a sparse, longer-range sub-network with 500 stronger connections. Connections for both sets of synapses were sampled with connection probability decaying exponentially with distance. Simulating whisker stimulus, we adjusted the initial state and input of the center 1 mm^2^-diameter circle to produce a sparse traveling wave of spreading activation similarly to the original publication. We altered the original model by normalizing synaptic strength by the average number of incoming connections and by basing connection probability on Euclidean distance rather than Manhattan distance. Moreover, because parameters such as membrane capacitance and synaptic weights were not provided in the source publication, we assigned values that approximated their results. For exact values, see the code repository.

We configured Cleo to simulate an experiment with an “optrode” (a combined electrode and optic fiber) to trigger inhibitory optogenetic stimulation when recorded multi-unit activity reached one or more spikes over the previous 0.5 ms sampling period (see Fig. 8a). To compensate for the unrealistically sparse network, we used a larger-than-default *r*_noise floor_ = 200 µm and did not simulate collisions. We simulated an inhibitory opsin with the previously described simple opsin model to accommodate the neuron model’s normalized, non-biophysical parameters and adjusted the strength through trial and error to a level sufficient to suppress activity around the optic fiber. To assess the effect of control latency, we also simulated the same experiment with an added 3 ms delay. The model was run for 15 ms of simulated time.

**Figure 8:**
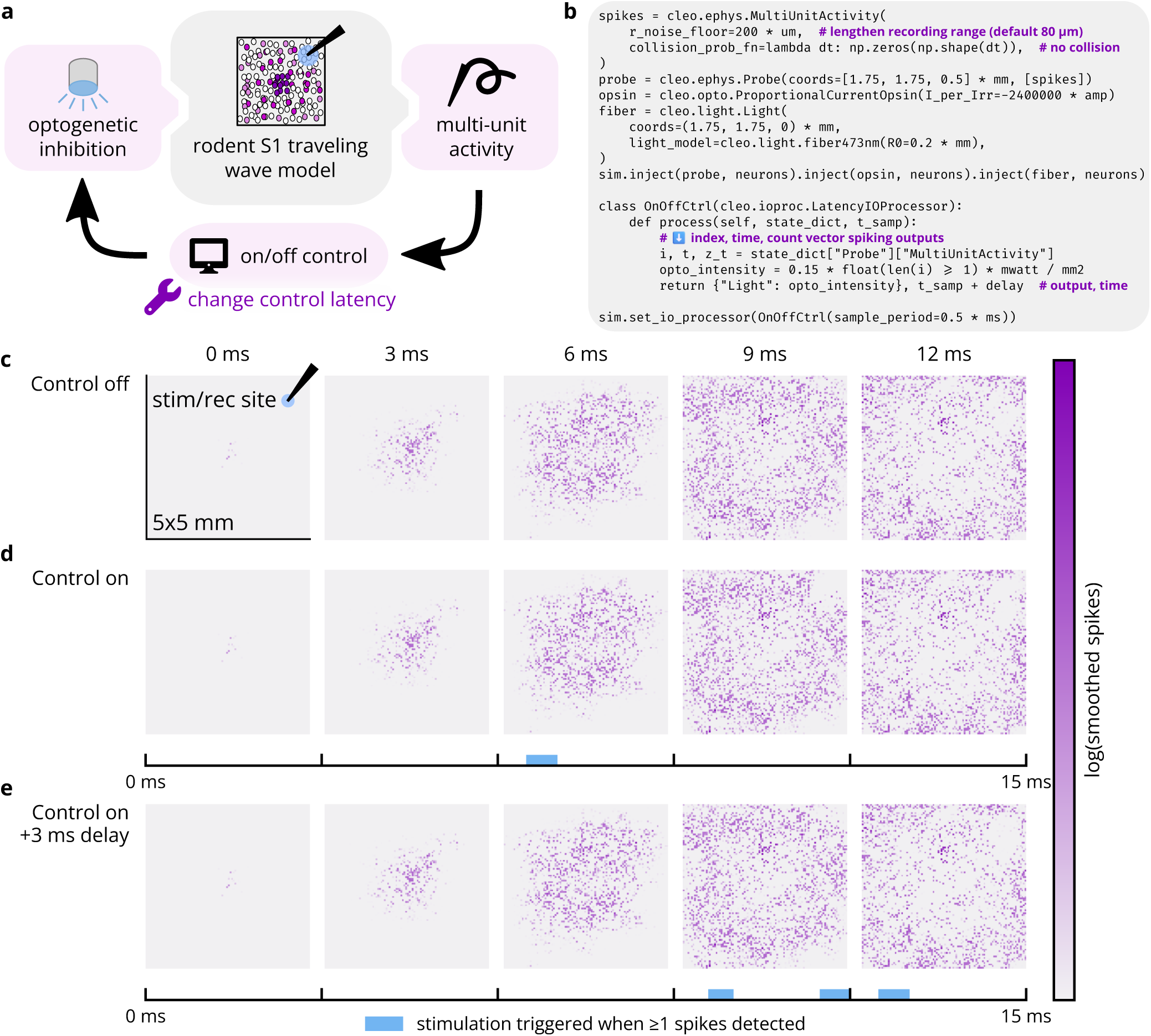
Cleo can simulate closed-loop inhibition of a whisker stimulation-evoked traveling wave. (A) Schematic of simulated experimental setup. The model consists of a 5 mm × 5 mm cortical area and optrode. The center 1 mm^2^-diameter circle of neurons is strongly stimulated, initiating a traveling wave of activity radiating outward. When sufficient spiking is detected at the electrode, an optical stimulus activating an inhibitory opsin is triggered. The experiment is simulated with and without closed-loop latency, demonstrating how Cleo can help determine requirements for real-time control. (B) Minimal code sample to configure the non-model components of the experiment. (C) Spatial spiking rasters over time. Each pixel represents the firing rate of a neuron, smoothed with a Gaussian kernel of 0.8 ms. (D) Top: Results of another simulation as in *C*, but with closed-loop inhibition. Neural activity is disrupted by the optogenetic stimulus in the neighborhood of the optrode. Bottom: Photostimulation over time. (E) Same as *D*, but with 3 ms latency introduced into the control loop. This latency prevents the controller from rejecting the traveling wave as it first enters the vicinity of the optrode (∼9 ms).

As seen in Fig. 8c,d, the optogenetic inhibition suppresses neural activity, effectively quenching the traveling wave in the region around the optrode. As expected, delay in the control loop prevents effective suppression of the traveling wave as it first reaches the optrode (see Fig. 8e). This demonstrates the use of Cleo in simulating basic “reactive” or “event-triggered” control where a predetermined stimulus is presented in response to a detected feature in the electrophysiology recording. In general, this sort of closed-loop control might be used to either inhibit (Ego-Stengel and Wilson, 2010; Krook-Magnuson et al., 2013) or amplify said feature. In this case, while constant inhibition could have achieved a similar effect, it would have posed a stronger intervention, increasing the likelihood of unnatural results. This prospective experiment also shows how Cleo can easily interface even with highly abstracted spiking neuron models.

#### 3.3.2. Prospective experiment 2: Clamping firing rate to disrupt plasticity in V1

Feedback control promises the ability to more tightly control variables of interest, enabling stronger causal conclusions about their downstream effects. In this prospective experiment, for example, we demonstrate how a closed-loop controller simplifies obtaining a consistent, desired firing rate of a subset of neurons in a primary visual cortex (V1) layer 2/3 plasticity model (Wilmes and Clopath, 2019), with the end of analyzing the effect on synaptic weight changes. A Brian 2 implementation of the model was publicly available on ModelDB (McDougal et al., 2017) and required only the minor modification of assigning coordinates to neurons (random locations in a cylinder of radius 84 µm and height 200 µm, the radius estimated using density data for mouse V1 from Herculano-Houzel et al. (2013)). This model features a variety of neuron subtypes, including pools of vasoactive intestinal peptide-expressing (VIP), somatostatin-expressing (SST), parvalbumin-expressing (PV), pyramidal (PC) cells, and L4 feedforward neurons (FF). The network is defined with inhibitory connections VIP-SST, SST-PV, SST-PC, and PV-PC; recurrent excitatory connections PC-PV and PC-PC (see Fig. 9a); and FF-PC and FF-SST feedforward synapses and gap junctions. A brief period (24.5 seconds) of top-down reward input to VIP neurons is sufficient to cause substantial changes to neural weights in a longer post-reward period. This is due to increased VIP activity inhibiting SST, disinhibiting PV, and strengthening a subset of SST-PV synapses such that a subset of PC cells are disinhibited (via PV) during a subsequent refinement phase. Thus, we concluded that disrupting PV activity during the reward period should be sufficient to disrupt plasticity in the PC connections. The model included a total of 694 neurons and 403,249 synapses, simulated over 137 seconds.

**Figure 9:**
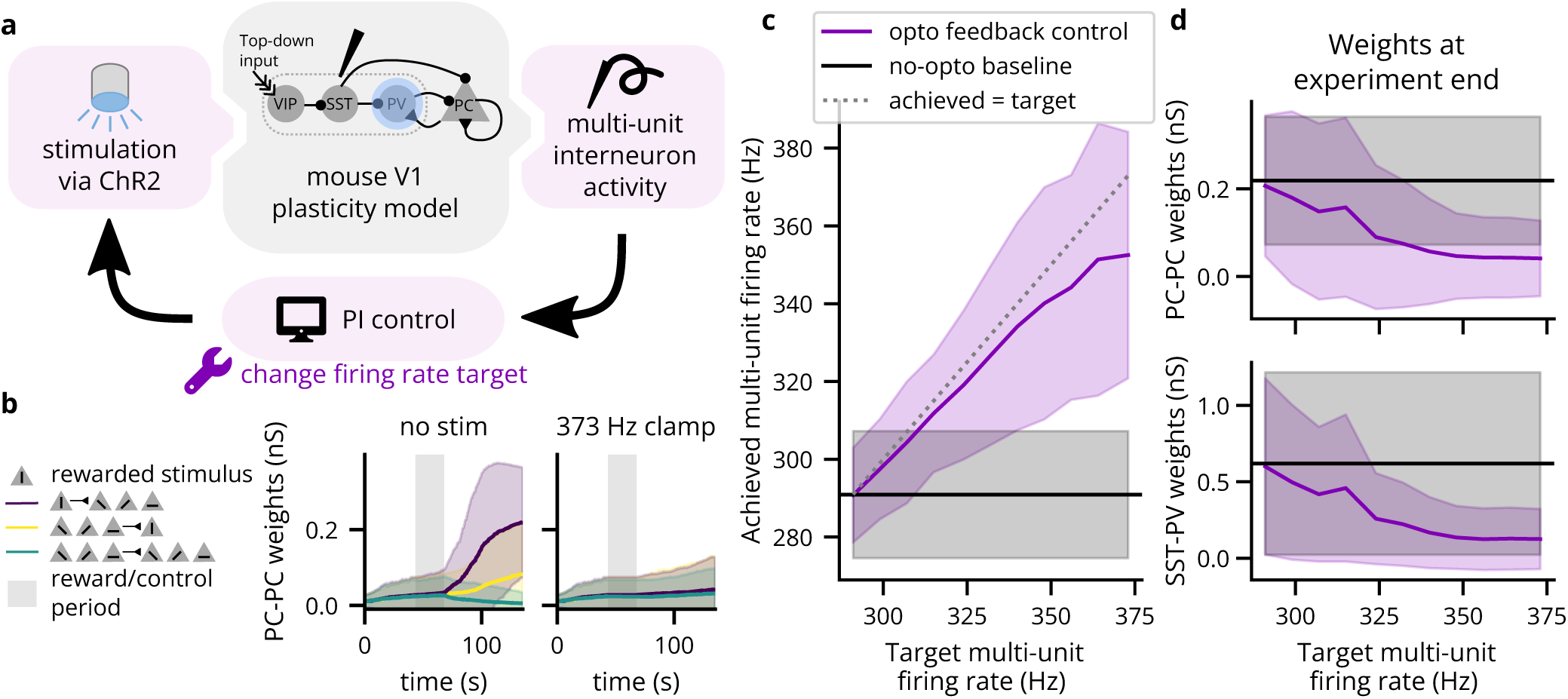
A Cleo simulation of optogenetic feedback control, clamping interneuron firing rate to disrupt top-down visual plasticity. (A) Schematic of the experimental setup. A model including simulated VIP, SST, PV, and PC neurons (Wilmes and Clopath, 2019) was perturbed via optogenetic feedback control. The PI controller set light intensity targeting PV interneurons transfected with ChR2. The experiment was repeated with different target firing rates. (B) The neural weights across time for PC-PC connections. Neurons are grouped by which stimulus they were selective for, where the vertical stimulus was rewarded. PC-PC connection weights from neurons selective to the rewarded stimulus (S) to nonselective neurons (NS) are shown in purple, NS-S in yellow, and NS-NS in green. Solid lines indicate the mean and shading indicates ±2 standard deviations. Top-down reward period is indicated by gray shading. Weights over time without (with) optogenetic control of firing rate are shown on the left (right). (C) Actual multi-unit, reward period firing rates for various targets. Solid lines/shading as in *B*. (D) The weights at simulation end (*t* = 126 s) for PC-PC, S-NS connections (top) and for reward-selective SST-PV connections (bottom). Solid lines/shading as in *B*.

We used Cleo to model an electrode recording multi-unit inhibitory activity (using Cleo’s default MultiUnitActivity parameters, with the exception of *r*_noise floor_ = 105 µm), simulating the scenario where the cell type of incoming spikes is identified in real time based on their waveform. To establish a baseline, we measured the mean and standard deviation of the detected firing rate during the reward period. Based on these results, we then ran control simulations with target reward period firing rate ranging from the mean to 10 standard deviations above the mean. We followed the methods in (Bolus et al., 2018), setting the light intensity in real time via a proportional-integral (PI) controller as implemented in the cleo.ioproc module. We used integral and proportional gains *K_i_* = 0.003 mW*/*mm^2^*/*spikes and *K_p_* = 0.005 mW*/*mm^2^*/*Hz. Firing rate was estimated via an exponential filter with time constant *τ* = 1 s.

PI control modulated firing rates in predictable ways that agreed with the goals of the experiment. Fig. 9b demonstrates how in the baseline simulation, reward-selective PC-PC connections are strengthened during the refinement period, whereas they remain relatively flat when elevating reward-period interneuron firing rate. Fig. 9c confirms that the PI controller successfully modulates reward-period interneuron firing rates, although with greater difficulty for higher targets. As expected, this attenuation of plasticity can be seen in both PC-PC and SST-PV connections, and this attenuation increases with the target firing rate (see Fig. 9d). While this sort of experiment is certainly feasible without feedback control, the open-loop alternative to attain a given reference firing rate would be the careful and potentially time-consuming titration of stimulation levels. In this way, Cleo has demonstrated a nominal prototype of an experiment where closed-loop optogenetic control can potentially be used to help elucidate the causal connection between components and phenomena of a network. This also demonstrates Cleo’s built-in PI control algorithms which provide users with an easy point of entry to feedback control.

#### 3.3.3. Prospective experiment 3: Evoking SWRs in the hippocampus

To demonstrate Cleo’s capabilities to simulate optimal feedback control of LFP, we interfaced Cleo with an anatomically informed model of the hippocampus previously described (Aussel et al., 2018, 2022)—the same used in Sec. 3.2.1 (see Fig. 10a,b). Denser than the epileptic version of the model, the healthy network features 33,200 neurons and ∼2.2 × 10^6^ synaptic connections. When a sustained external current is delivered to the entorhinal cortex to simulate the slow waves of non-REM sleep, the model produces a sharp wave-ripple (SWR)-like pattern of LFPs, as approximated by summed synaptic currents. Our goal was to evoke a SWR using optogenetics in the absence of this strong square-wave input, illustrating how feedback control can reproduce a signal of interest at arbitrary times. Moreover, feedback control replaces a design and calibration process with model fit and controller tuning, producing a stimulation waveform that need not conform to a basic shape. In contrast, various experimenters have used rectangular, trapezoidal, and ramping pulses to optogenetically induce SWR-like oscillations *in vivo* that do not fully resemble spontaneous SWRs, apparently manually calibrating the intensity (Stark et al., 2014; Oliva et al., 2018; Tingley et al., 2021).

**Figure 10:**
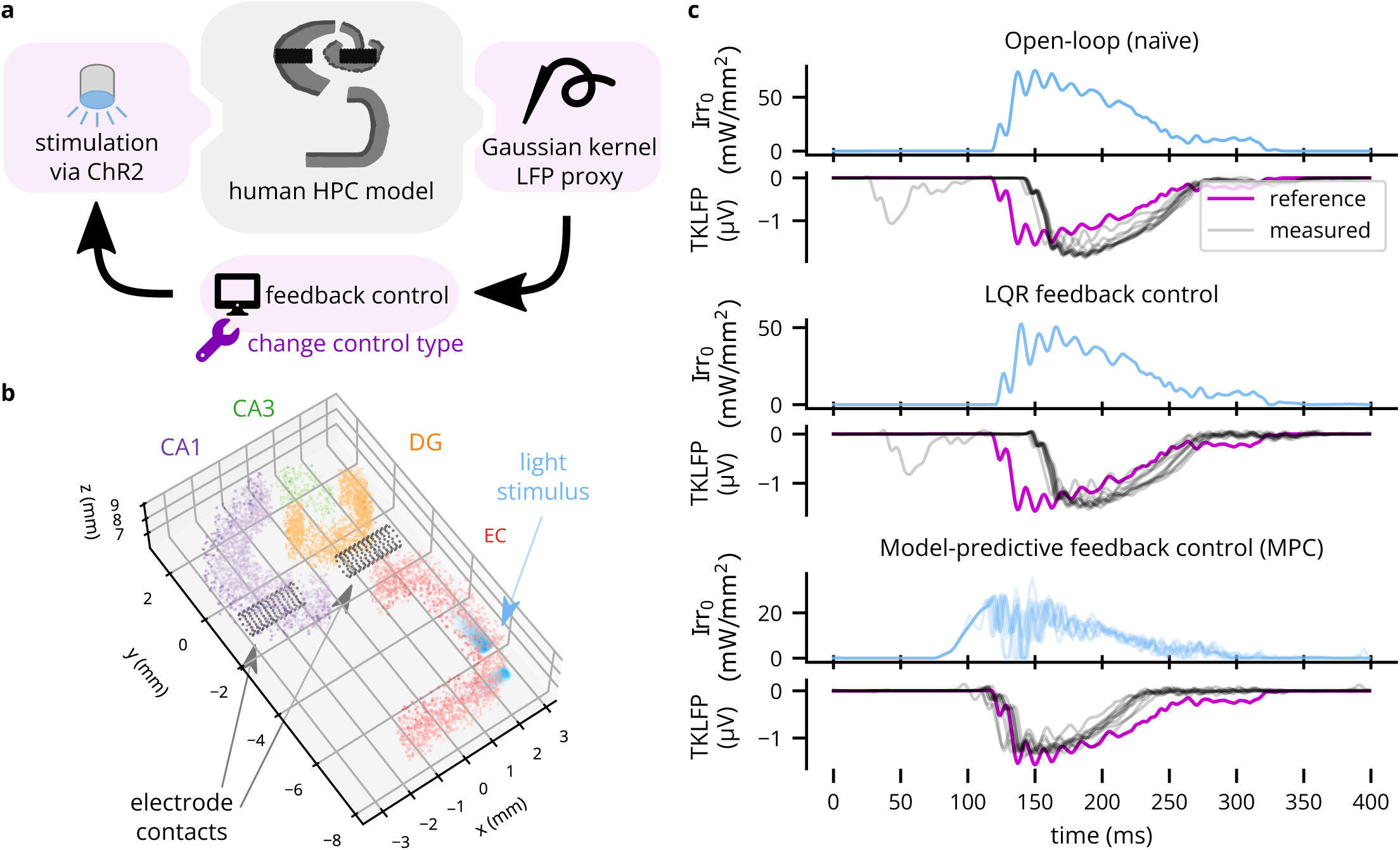
An example application of Cleo using optimal feedback control to follow a time-varying waveform. (A) Schematic of the simulated experiment setup. TKLFP is recorded from the anatomical hippocampus model by Aussel et al. (2018) and is fed into a feedback controller governing ChR2 photostimulation. Different control methods are simulated to show Cleo’s utility in assessing different candidate stimulation protocols. (B) A 2.5 mm-thick slice of the 15 mm-tall model is shown. The model consists of four regions, entorhinal cortex (EC), dentate gyrus (DG), CA3, and CA1. Electrode contacts are represented as black dots and are in the same location as in the original model. Two light sources are shown in EC. Nine other such pairs (for a total of 20 light sources) not pictured here were spaced regularly parallel to the z axis. (C) Results are shown for ten trials each of three stimulation paradigms: naïve open-loop, LQR, and MPC. Input Irr_0_ is the light intensity at the tip of each optic fiber.

In the Cleo simulation, we placed simulated electrode contacts at the same locations as in the original model and used them to record LFP using the TKLFP approximation (Telenczuk et al., 2020). We then inserted optic fibers along the 15 mm model length and injected Gaussian process noise into the external current driving the model to create trial-to-trial variability. We illustrate three stimulation paradigms: naïve open-loop (consisting of a mirrored, rectified version of the reference signal), linear quadratic regulator (LQR), and model-predictive control (MPC). The results demonstrate that Cleo can be used to simulate complex experimental scenarios with multiple recording and stimulation interfaces, and a variety of stimulation protocols can be prototyped on the same model with relative ease. In this case, the simulated response to stimulation is quite stereotypical, creating little meaningful trial-to-trial variation for the advantages of LQR over open-loop control to become apparent (see Fig. 10c). MPC, however, produces a notably earlier response than LQR since it is able to “look ahead”.

## 4. Discussion

Here we have presented Cleo, a Python package designed as a testbed for bridging point neuron spiking network models and experiments for mesoscale neuroscience. As the sole publicly available tool for simulating delayed closed-loop control, two-photon optogenetics, and multi-opsin/wavelength crosstalk, Cleo excels in consolidating various niche models into one adaptable platform, sparing researchers the need to understand and implement them on a case-by-case basis into their SNN simulations. By thus simulating the experimental apparatus, Cleo can facilitate the process of model informing experiment and experiment informing model, which is a bidirectional research paradigm often advocated as providing the richest potential understanding of brain function.

In the design and prototyping phase, Cleo can help answer beforehand questions such as whether an experiment is feasible (Antolik et al., 2021), which opsin(s) or indicator(s) to use, what cells to target, where to record, or what closed-loop control algorithms perform adequately and with tolerable latency. By simulating the messy side effects of each choice, Cleo can help narrow down a number of suboptimal alternatives and make trade-offs between competing constraints. When a sufficiently realistic model for the studied system does not exist, multiple models representing possible variations in connectivity, parameters, or mechanisms could be used to cast light on which experimental configurations work best across hypothetical models. Indeed, the desired experiment in this case could be one that best adjudicates between these hypotheses (Willats, 2022). Other potential applications of Cleo include facilitating the reverse process of experiment informing model (Gutzen et al., 2018; Trensch et al., 2018; Gerkin et al., 2019; Appukuttan et al., 2022) and guiding methods engineering, including control algorithms, calcium indicators, and opsins, for example.

A primary motivation for developing Cleo is to accelerate the development of closed-loop optogenetic control (CLOC), which may enable stronger causal hypothesis testing. Neuroscientists have identified many network– and circuit-level variables and phenomena in search of interpretable explanations of brain activity—controlling these features is thus a natural application of CLOC. Examples of these features include cell type-specific activity; the type (Cole and Voytek, 2017), frequency (Saleem et al., 2017), amplitude (Saleem et al., 2017), spike coherence (Buffalo et al., 2011; Buschman et al., 2012) and interactions (Aru et al., 2015; Zhang et al., 2019) of oscillatory patterns; discrete phenomena such as bursts, sharp wave-ripples, oscillatory bursts (Akam and Kullmann, 2014; Lundqvist et al., 2016; Karvat et al., 2020), traveling waves (Muller et al., 2018), or sleep spindles (Fernandez and Lüthi, 2020); and latent states describing neural dynamics (Churchland et al., 2012; Vyas et al., 2020; Shenoy and Kao, 2021; Peixoto et al., 2021), including those most relevant to behavior (Sani et al., 2024; Hurwitz et al., 2021).

While some of these targets lend themselves easily to CLOC, others require continued innovation in interfacing technology. Specifically, stimulation technologies have been much more limited in their degrees of freedom than modern recording technology, and thus unlikely to sufficiently control many variables of interest. For this reason, the development of multi-channel micro-LED/optrode devices (Dufour and Koninck, 2015; Kwon et al., 2015; Welkenhuysen et al., 2016; Kampasi et al., 2018; Wang et al., 2018; McAlinden et al., 2019; Mao et al., 2019, 2021; Ohta et al., 2021; Antolik et al., 2021; Jeon et al., 2021; Kathe et al., 2022; Eriksson et al., 2022) and holographic optogenetic stimulation (Rickgauer et al., 2014; Packer et al., 2015; Ronzitti et al., 2017; Adesnik and Abdeladim, 2021; Yang and Yuste, 2021; Sridharan et al., 2022) are of particular interest. Crucially, Cleo will enable rigorous investigation of both proposed specific technologies as well as general technological capabilities to guide new interface design.

While Cleo was designed to facilitate and accelerate the simulation of complex experiments as much as possible, it has several limitations. First, while Brian and Cleo have the flexibility to accommodate a wide variety of models, alternative tools and methods—adapted as necessary to simulate the experimental interface—may be better suited for larger spatiotemporal scales (Sanzleon et al., 2013; Neymotin et al., 2020; Dai et al., 2020), higher levels of abstraction (Macke et al., 2011; Yamins et al., 2014; Pandarinath et al., 2018), and greater biophysical detail (Hines and Carnevale, 1997; Gratiy et al., 2018; Dai et al., 2020; Dura-Bernal et al., 2019). A second limitation is speed; Cleo cannot take advantage of Brian’s full compilation capabilities due to the arbitrary Python code that enables flexible recording/stimulation modules and closed-loop control.

Perhaps the biggest limitation is that the user must work to interface their model with Cleo, which could range from the simple task of assigning neuron coordinates to the considerable effort of re-implementing the model entirely with Brian, if not already a Brian model. Conversion from other simulators may be possible via NeuroML (Cannon et al., 2014), but Brian’s and NeuroML’s converter functionality is limited. Ideally, an experiment simulation testbed would flexibly support multiple simulation backends, as PyNN has provided for SNNs (Davison et al., 2009). To do so in a native, computationally efficient way would require significant work, using the idiosyncrasies of each simulator to implement features they were not designed for (e.g., opsins, lights, and calcium indicators), as we have done for Brian. A future collaborative effort extending a multi-simulator framework such as PyNN for this purpose may be worth the investment if there is enough community interest in expanding the open-source SNN experiment simulation toolbox.

Cleo is open-source and can be installed from the Python Package Index under the name “cleosim”. The code can is hosted on GitHub at https://github.com/siplab-gt/cleo, where we invite users to submit feature requests, bug reports, pull requests, etc. Documentation, including an overview, tutorials, and API reference, can be found at https://cleosim.readthedocs.io. Future development of Cleo is relatively straightforward given Cleo’s modular structure. We anticipate future development to meet community needs may include simulation of different levels of abstraction (e.g., forward modeling of extracellular potentials (Pettersen et al., 2012; Hagen et al., 2018) for multi-compartment models or additional light propagation profiles (Liu et al., 2015)), additional/improved recording and stimulation modalities (e.g., photoelectric artifacts, voltage imaging, two-photon imaging/optogenetics crosstalk, electrical micro-stimulation, or an expanded selection of opsins and sensors), or support for heterogeneous sampling rates to capture scenarios such as when imaging is slower than electrode recording (as in the real-time processing software Bonsai (Lopes et al., 2015), for example).

## 5. Author contributions

KAJ designed and developed the software, executed the validation experiments and the third prospective experiment, and wrote most of the manuscript and the documentation. NAC identified, simulated, and wrote the results for the first two prospective experiments, as well as helped maintain code. ZCM implemented model-predictive control for the third prospective experiment. CJR, JEM, and AAW provided direction and consultation and CJR additionally edited the manuscript. ASC and JEM gave direction and feedback on methods for modeling two-photon calcium imaging, and JEM additionally advised 2P optogenetics modeling.

## 6. Acknowledgments

This work was supported by the National Institutes of Health (NIH) NINDS CRCNS grant R01NS115327. KAJ was additionally supported by the Georgia Tech/Emory NIH/NIBIB Training Program in Computational Neural Engineering (T32EB025816). JEM is funded by a Career Award at the Scientific Interface from the Burroughs Wellcome Fund, a Sloan Foundation Fellowship, and a Fellowship from the David and Lucille Packard Foundation. We thank Michael Bolus, Cameron McIntyre, Thomas Foutz, Sukhdev Roy, Hillel Adesnik, Andrew Davison, Marcel Stimberg, Amélie Aussel, Jon Newman, Katharina Wilmes, Alberto Mazzoni, Gaute Einevoll, Márton Rózsa, Alain Destexhe, and Bartosz Teleńczuk for responding to inquiries regarding models, data, and code presented in their publications. We also recognize Tobias Niebur for contributing code used in Sec. 3.3.1 and Olivia Klemmer, Chuyu (Alissa) Wang, and Aarav Shah for contributing to the wslfp package.

## 7. Supplementary Methods

### 7.1. Multi-wavelength sensitivity

#### 7.1.1. Light superposition

Here we describe how we model dynamics for light of two different wavelengths, given photon flux *ϕ_λ_*_1_ and *ϕ_λ_*_2_. Bansal et al. (2020b) use a weighted sum of activation functions

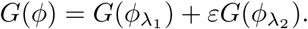

However, these are sublinear functions, so adding them results in exaggerated activation. In the extreme case, imagine two light sources that are just 1 nm apart in wavelength, with *ε* = 1. Thus:

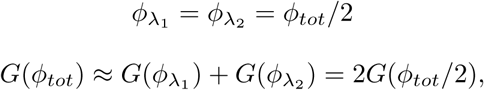

which contradicts what we expect for sublinear *G*:

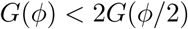

A more accurate approach instead assumes the following

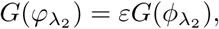

where *φ_λ_*_2_ is the standard-wavelength equivalent (“effective flux”) of *ϕ_λ_*_2_.

Demonstrating with *G_a_*_1_ of the four-state model (Evans et al., 2016), we solve for *φ* in terms of *ϕ* and *ε*:

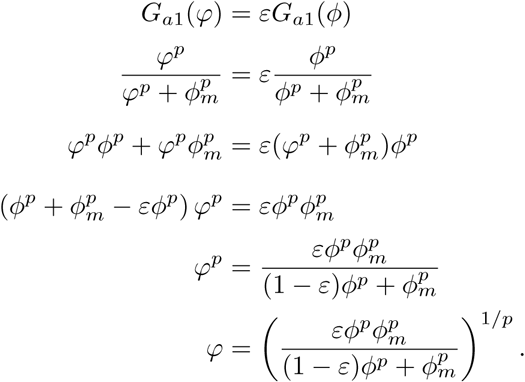

We then compute our activation functions as

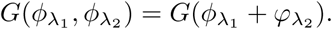

Unfortunately, this doesn’t yield a simple constant conversion factor. However, if we approximate *G* as linear, we can use a weighted sum of fluxes:

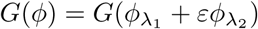

Plotting *G* for multiple opsins shows this linear approximation does yield a lower activation curve than the Bansal *et al*. approach and is qualitatively close to the true *φ* derived above. See Fig. S6 for a comparison of the three methods.

#### 7.1.2. Action spectrum normalization

Some action spectra are measured with equal power density/pulse width across wavelengths, while others are reported with equal photon flux. We store ours as equal power density spectra since they seem to be more common and allow for the opsin model to use both accurate power density and photon flux values. We use *ε* to represent sensitivity relative to the peak-sensitivity wavelength, *ε_ϕ_* and *ε_P_* representing the equal photon flux and power density versions, respectively.

For a given wavelength *λ*,

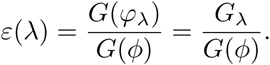

We let *G_λ_* represent the response at wavelength *λ*, while *G*(*ϕ*) represents the response at the peak wavelength for the same flux *ϕ*. We will assume *G* is a linear function, as above, using *C* to represent a constant:

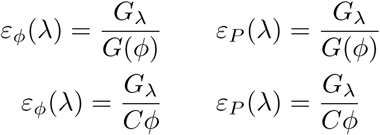

Then we make either photon flux or irradiance (power density) constant:

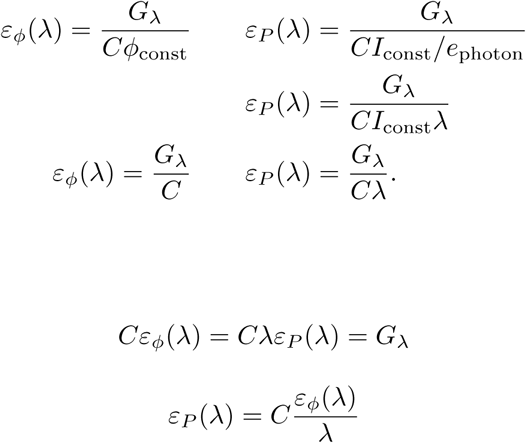

Thus, we can convert from *ε_ϕ_*(*λ*) to *ε_P_* (*λ*) by dividing by *λ* and normalizing.

### 7.2. GECI convolution simulation as an ODE

Song *et al*. convolve the intracellular calcium trace [Ca^2+^] with a double exponential kernel to capture variable rise and decay times in the fluorescence signal. To simplify simulation (to not have to keep a buffer of past calcium values), we can represent this convolution as an ODE. Let *c*(*t*) and *b*(*t*) be the free and bound calcium concentrations and *h*(*t*) be the kernel function

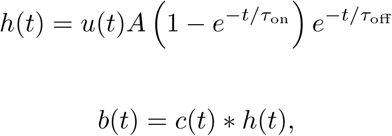

where *u*(*t*) is the unit step function, included to ensure the kernel is causal.

We can represent the convolution as multiplication in the Laplace domain:

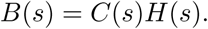

By expanding out *h*(*t*), we get functions that can easily be transformed into the Laplace domain. Let 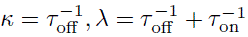 to simplify notation.

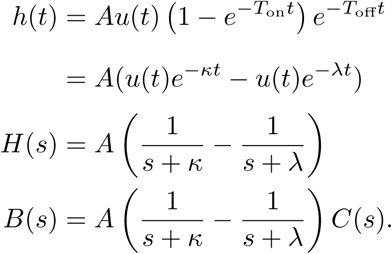

Now we get a common denominator and rearrange:

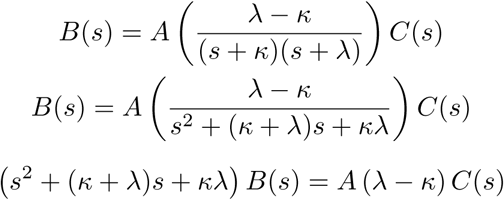

Now we use the *s*^2^ and *s* terms to convert to a second-order ODE, using the fact that the Laplace transform of *b*^′′^(*t*) and *b*^′^(*t*) are *s*^2^*B*(*s*) − *sb*(0^−^) − *b*(0^−^) and *sB*(*s*) − *b*(0^−^), respectively. Also, we assume that *b*(0) = *b*^′^(0) = 0 to avoid undefined *δ*(*t*) and *δ*^′^(*t*) terms after the inverse Laplace transform; this appears to have only a minor effect.

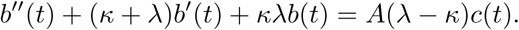

Rearranging to a first-order ODE system by introducing *β*(*t*) = *b*^′^(*t*), we get

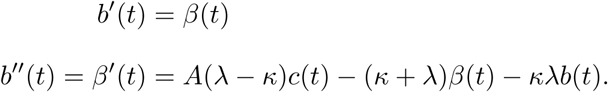

Special thanks to DinosaurEgg on Math Stack Exchange for helping solve this problem.

### 7.3. Probabilistic spike detection details

To approximate multi-unit and sorted spike recording without simulating full extracellular action potential (EAP) waveforms, Cleo takes ground-truth spikes returned by the Brian simulator, makes noisy measurements of their amplitudes alone, and stochastically determines which to report as recorded on the probe. The recorded amplitude of an average spike *A_µs_* is a function of *r*, the distance between the neuron and the electrode (see Fig. 3a). This function is parameterized by *r*_noise floor_, the distance at which signal-to-noise ratio (SNR) is 1, i.e., *A_µs_*(*r*_noise floor_) ≜ *σ_b_* ≜ 1, where *σ_b_* is the standard deviation of background (electrical and biological) noise. The default *r*_noise floor_ is 80 µm, as reported by Cohen and Miles (Cohen and Miles, 2000), which is consistent with other literature stating the measured amplitude drops to near 0 between 140 and 300 µm (Henze et al., 2000; Somogyvári et al., 2005). Cleaner/noisier recordings can be achieved by lengthening/shortening *r*_noise floor_.

The shape of the amplitude profile is defined by 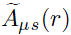, which can take any monotonically decreasing form, but is 1*/r*^2^ by default. This follows detailed simulations by Pettersen and Einevoll (Pettersen and Einevoll, 2008) that show *A_µs_*decreasing with 1*/r^n^*, where *n* varies between 2 and 3 depending on cell type and distance. The raw 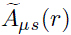 function is shifted by a small distance *r*_0_ = 5 µm to avoid division by 0 and scaled such that *A_µs_*(*r*_noise floor_) = *σ_b_*:

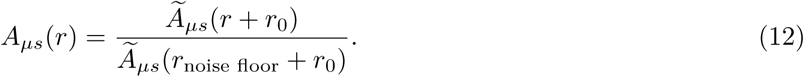

Variability in measured EAP amplitudes comes from the aforementioned background noise as well as variability in spike amplitude. The former is simulated as white noise in the spiking band, independent across channels, and scaled to have standard deviation *σ_b_*. To ensure the noise has consistent energy and spectral properties with different temporal resolutions, we produce a spiking band-limited noise waveform per channel. An alternative of a simple Gaussian sample every time step, for example, would produce more false positives the smaller the time step. Cleo produces this noise by filtering per-time step white noise with a 6th-order Butterworth filter with cutoff frequencies of 300 and min(3000, 0.45*f_s_*) Hz, the upper cutoff being adjusted to stay under the Nyquist frequency.

Intrinsic amplitude variability is modeled as a Gaussian distribution around the mean with standard deviation CV*_A_A_µs_*. CV*_A_* signifies the coefficient of variation and is 0.05 by default. This default was estimated from the median of 20 randomly selected reporter-positive mouse cells from the Allen Cell Types Database (all). For each cell, we selected the sweep with the most spikes and computed the coefficient of variation of spike amplitudes, where amplitudes were taken to be the difference between the peak and threshold voltages for each spike.

Combining variation from these two sources, the variance in measured amplitude across spikes is thus 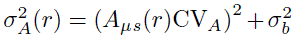. With *A_µs_*(*r*), *σ_A_*(*r*), and detection threshold *ϑ*, we can compute the probability of detecting a given spike from a neuron on a single channel at distance *r* as

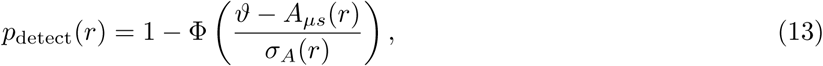

where Φ is the CDF of the standard normal distribution (see Fig. 3a).

In large simulations, it would be computationally expensive to sample threshold crossings and especially collisions for neurons that are distant from the electrode and/or each other. Thus, when a probe is injected into a neuron group, Cleo computes the multi-channel recall for each neuron as

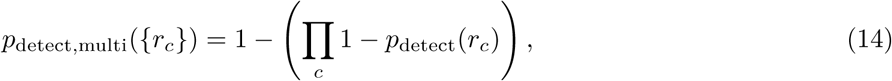

where *r_c_* is the distance from the neuron to channel *c*. The probe then subscribes only to neurons with *p*_detect,multi_ *>* 0.001, thus including fairly low-SNR neurons but ruling out the very distant. Note that while noise is independent across channels, the intrinsic spike amplitude is shared. The independence assumption in Equation 14 is thus an approximation that saves computational effort by removing the correlation cross-terms across channels.

Candidate threshold crossings are identified from spikes and non-spike noise values that exceed *ϑ*, by default 4*σ_b_*. To efficiently consider all spikes on a given channel in parallel and support varying temporal resolution, they are assessed independently despite the fact that in a real experiment there is only one measurement on a channel at a given time. I.e., in our method two subthreshold measurements do not combine to produce a threshold crossing. While this approach may seem unrealistic, it avoids another unrealistic scenario where many distant spikes combine to regularly exceed the threshold. This also supports varying temporal resolution in the sense that it allows us to use coarser temporal resolution without limiting how many “measurements” can be made at each time step per channel.

These threshold crossings then go through a “collision” sampling process modeling limitations in detecting overlapping spikes. This is parameterized by a collision probability function *p*_coll_(Δ*t*), where Δ*t* is the time difference between candidate threshold crossings. To capture the common practice of setting a minimum refractory period in MUA, the default for multi-unit activity is *p*_coll,MUA_ = **1**_{Δ_*_t<_*_1 ms}_, meaning that collision is certain for spikes within 1 ms of each other. Reflecting the behavior typical of many spike sorters (Garcia et al., 2022), the default for sorted spiking is an exponential decay function *p*_coll,SS_ = 0.2 exp(−Δ*t/*0.3 ms) (see Fig. 3b). Every spike is compared to previous spikes in the recent past and in the case of collision, the first one is kept (to preserve the causal nature of the simulation). For simultaneous spikes, the one with the highest amplitude is kept.

The resulting output is then processed to produce the final multi-unit or sorted output (for an example, see Fig. 3c,d). Multi-unit activity reports every spike detected by every channel, without regard for the origin of the spike (though with the default collision function, there will never be more than one per time step per channel). Sorted spiking, on the other hand, reports all spikes detected on at least one channel, where each neuron is identified by a unique index. Only neurons with a SNR (defined as the peak *A_µs_/σ_A_* across channels) above a threshold (default 6*σ_b_*) are included in the sorted output, reminiscent of SNR filtering of spikes in some sorters (Yger et al., 2018) and analyses (Magland et al., 2020). While real-time spike sorting for closed-loop control is difficult in practice for large channel counts, this sorted spiking option could be used to emulate a more common workflow of isolating one or a few neurons to record spikes from in real time. Note that we simplify the sorting process in several ways: (1) by assuming perfect sorting, (2) by treating the detection process as an independent process per channel, not incorporating spatial patterns as spike sorters do, (3) by keeping the first spike in case of collisions, rather than throwing out both or misclassifying one, and (4) reporting no false positives, since threshold crossings from noise alone would very rarely match a template. As a consequence of the last point, the downside to lowering the detection threshold for sorted spikes is that false positives can increase spikes missed due to collisions.

### 7.4. Bayesian optimization of neuron models for light pulse characterizations

Neuron parameters were selected for AdEx and LIF neurons to match the firing rates of the source simulations in (Foutz et al., 2012) via Bayesian optimization. We used the Bayesian Optimization Python package (Nogueira, 2014; Gardner et al., 2014), primed with four parameter settings from the Neuronal Dynamics Textbook (Gerstner et al., 2014) and 20 random settings, followed by 200 iterations of optimization. The objective function was the sum of squared error between the target and simulated firing rates, totalled across all five conditions, i.e., 120%/75%, 120%/100%, 120%/150%, 100%/100%, and 140%/100% light intensity/expression level combinations. 100% light intensity was defined as that required to produce a single spike with a 5 ms pulse.

### 7.5. Prospective experiment 3

The reference signal was generated by delivering a 1 nA square wave input from 100 to 300 ms to entorhinal cortex using the original model’s *I*_ext_ term, without noise added. Training data was generated by running the system for 13 seconds with alternating on and off periods of length *T* ∼ N(200, 50) µs. During “on” intervals, *I*_ext_ ∼ |N (0, Irr_max_*/*3) |. Irr_max_ was 75 mW*/*mm^2^, which is described as an upper safety limit for 473 nm light delivery to the brain (Cardin et al., 2010). Gaussian process noise was generated with mean *µ* = 0.167 nA and using the exponentiated quadratic kernel

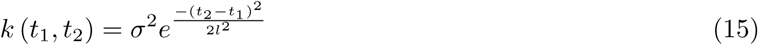

with *σ* = 0.083 nA*, l* = 30 ms and was added to input current *I*_ext_ in each control scenario. The parameters of light delivery were altered from the 473 nm-wavelength optic fiber defaults to allow for greater propagation—*K* and *S* were both divided by 10. The training data, sampled at 1 kHz, was fit using the ldsCtrlEst library’s SSID and EM fitting methods with latent dimensionality *n_x_* = 4 (Bolus et al., 2022). An SSID fit was performed first, later refined by EM. ldsCtrlEst’s Gaussian linear quadratic regulator (LQR) controller was used with a gain computed from *Q* = *C^T^ C, R* = 0.001 state and input penalties, with a simulated 3 ms of latency.

Model-predictive control (MPC) was implemented with a control horizon of 50 time steps, i.e., 50 ms. The standard, quadratic cost function utilized the same *Q, R* as LQR, and was optimized using the ‘cvxpy’ Python package (Diamond and Boyd, 2016). MPC was simulated with 6 ms latency.

## 8. Supplementary Figures

**Figure S1:**
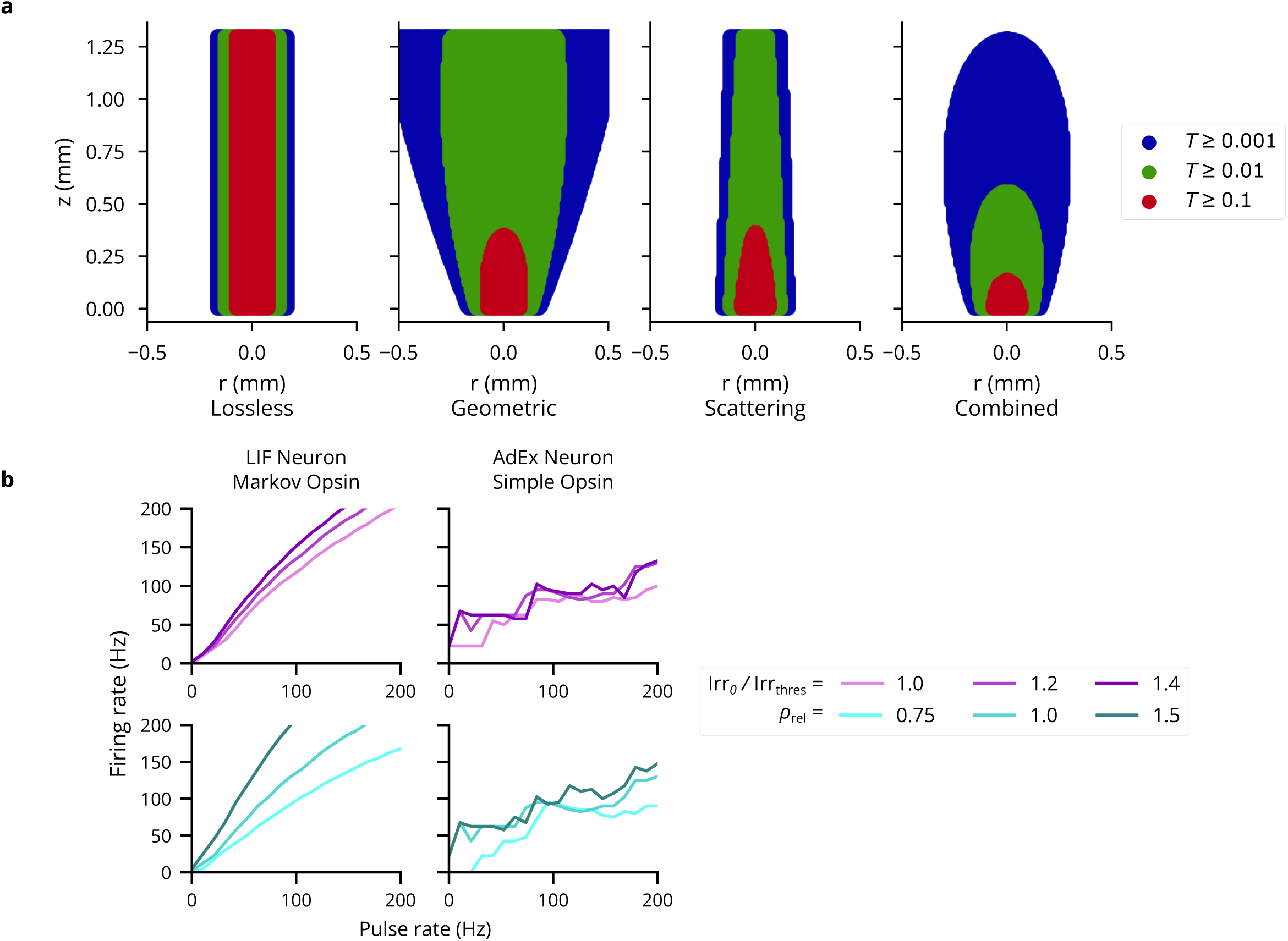
(A) Light transmittance *T* as a function of radius and axial distance from the optic fiber tip (cf. Figure 2a from (Foutz et al., 2012)). The contribution of the Gaussian distribution, cone-shaped light propagation, and scattering are depicted separately. (B) Firing rate-pulse rate relationship as in Fig. 4c, for more neuron model-opsin combinations, namely LIF neuron with four-state Markov opsin and AdEx with a proportional current opsin. 5 ms pulses are used as before, with irradiance and expression levels as shown in the legend.

**Figure S2:**
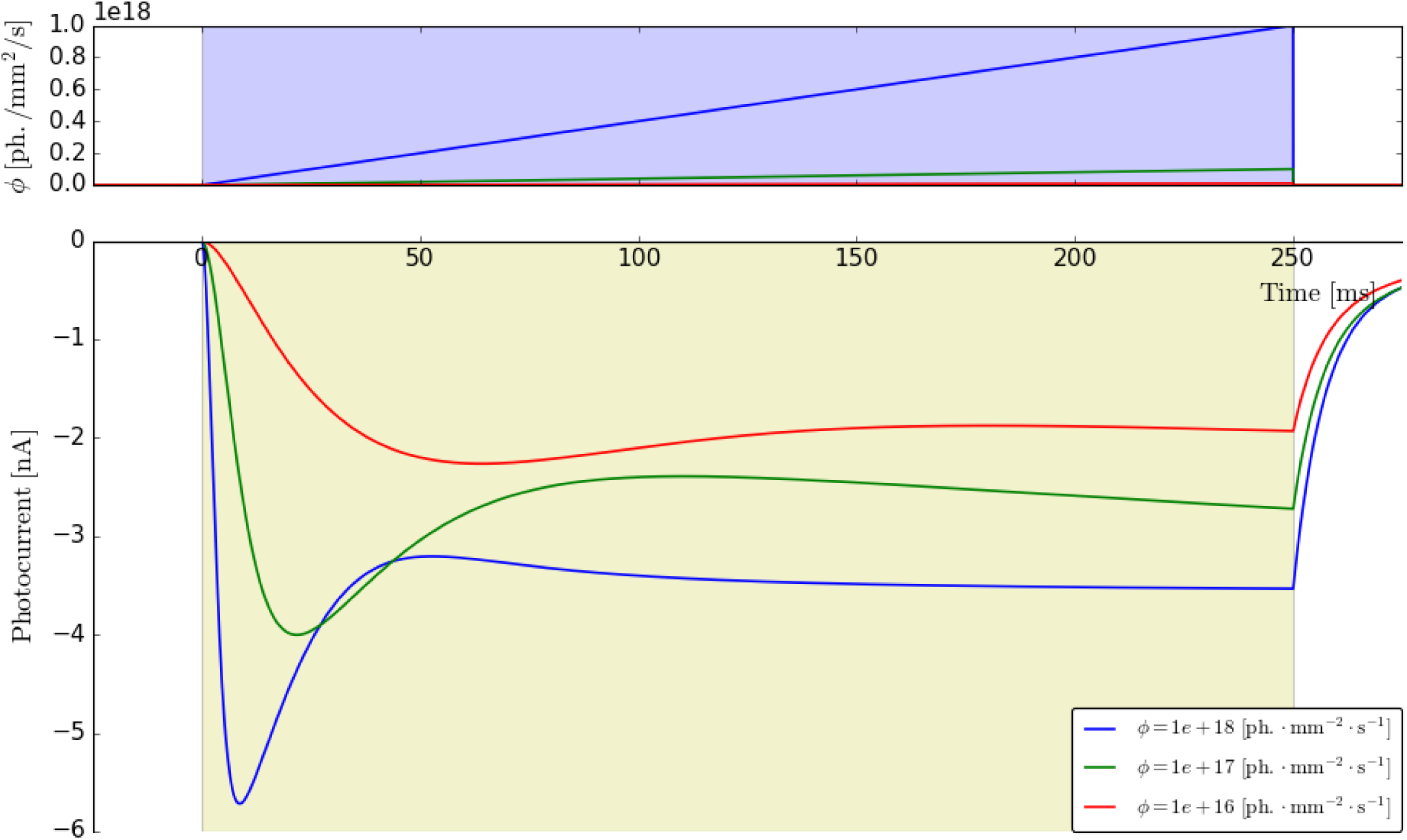
Output from the “ramp” protocol for the four-state ChR2 opsin model, produced by the PyRhO optogenetics simulation platform (Evans et al., 2016).

**Figure S3:**
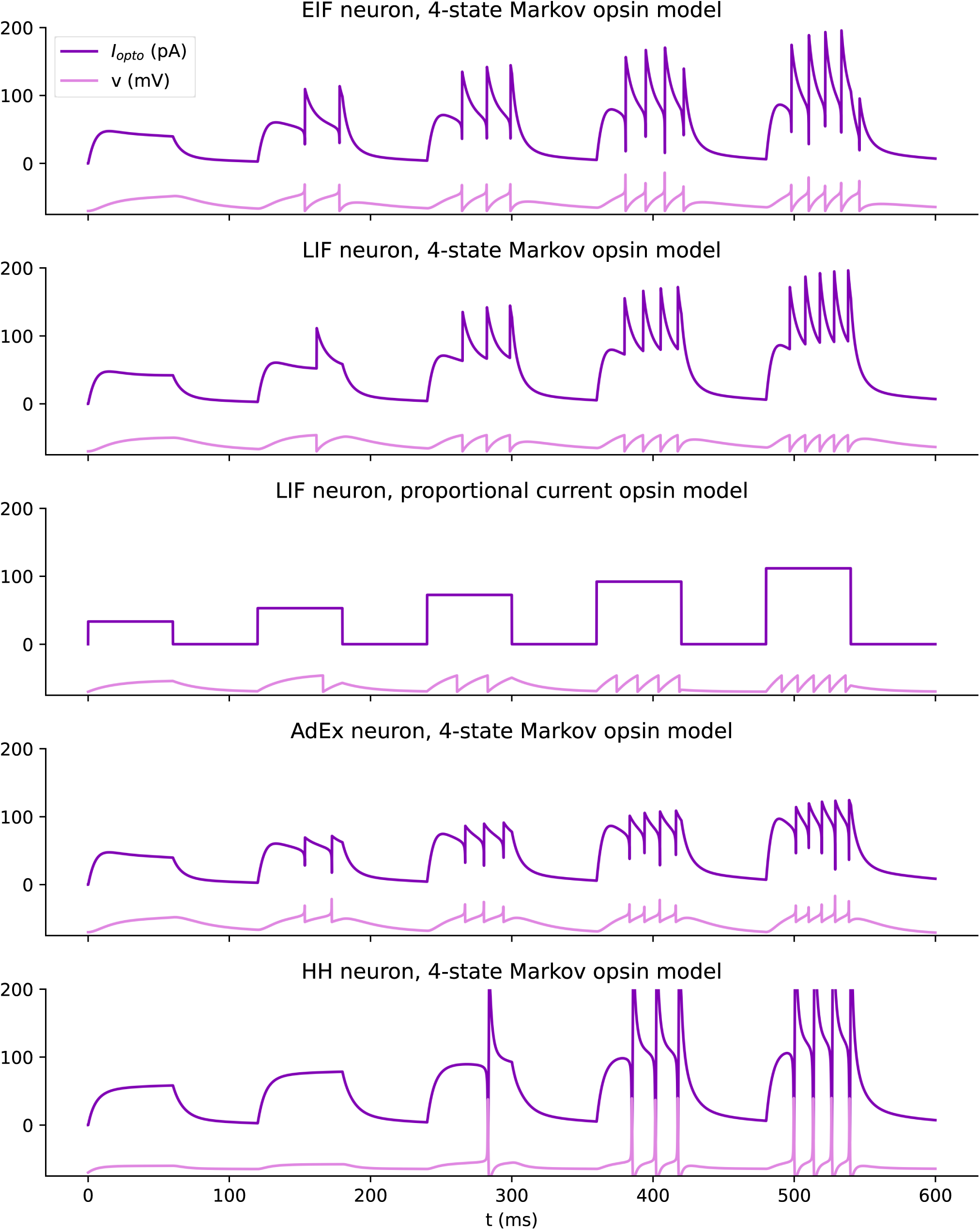
Responses of diverse opsin/neuron model combinations to light pulses of increasing intensity show qualitatively similar light-firing rate relationships.

**Figure S4:**
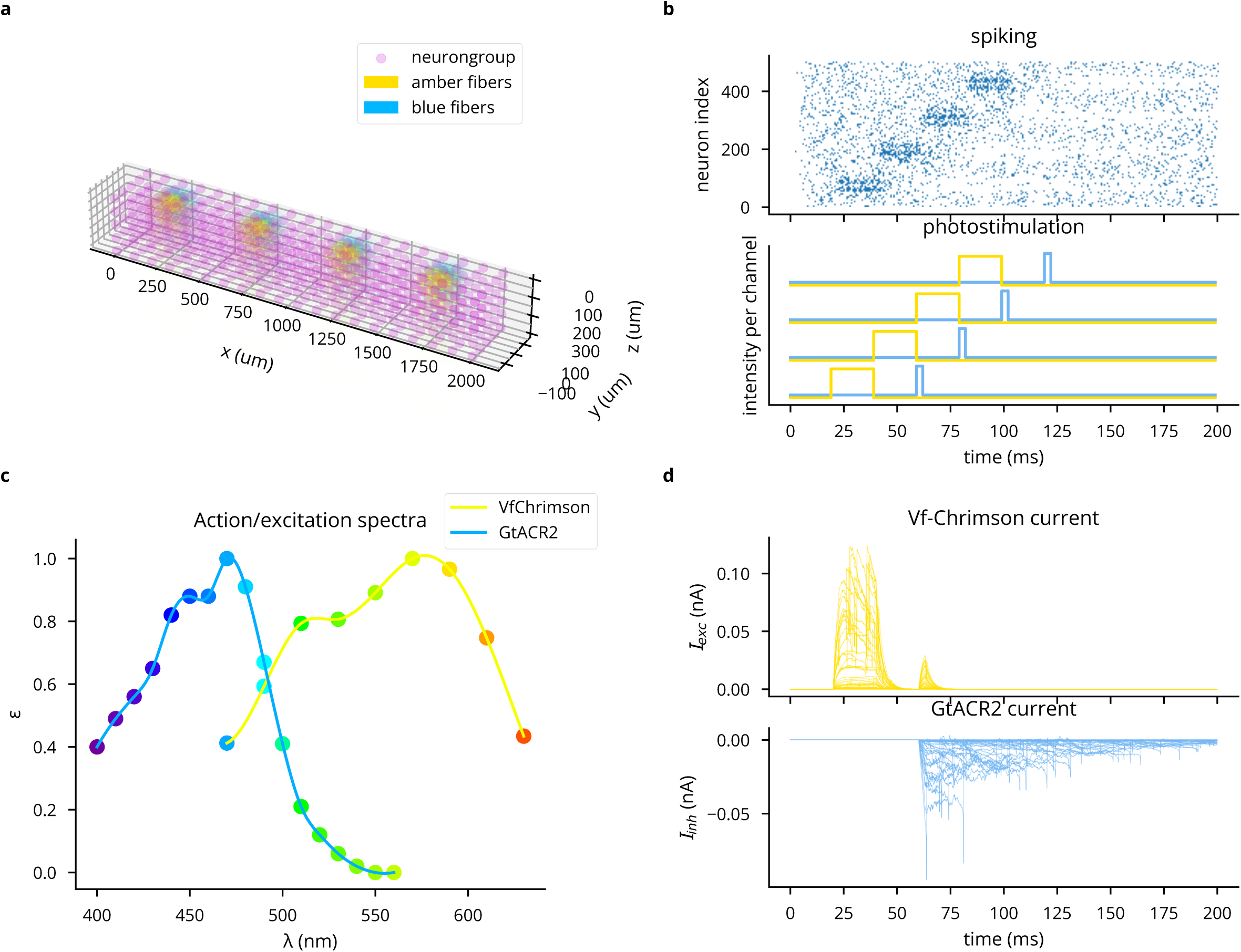
Demonstration of simulating multiple light sources, wavelengths, and opsins simultaneously. (A) 3D plot of network model and light sources. (B) Top: spike raster, where increasing neuron index correlates with increasing *x* coordinates. Bottom: Stimulation pattern for 473 and 590 nm light sources. (C) Action spectra of Vf-Chrimson and GtACR2, showing crosstalk of blue light on Vf-Chrimson. (D) Photocurrents for the first 50 neurons.

**Figure S5:**
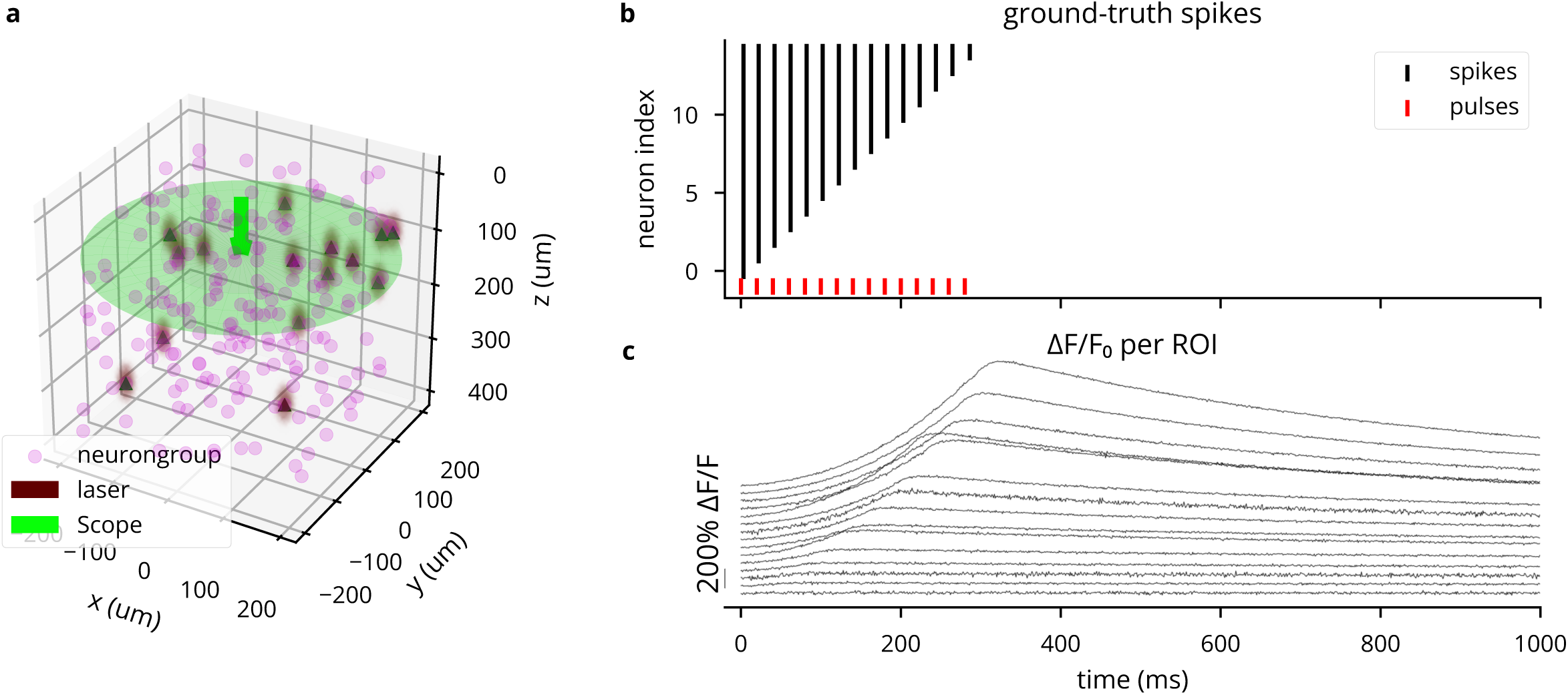
Simulation of two-photon calcium imaging using the GCaMP6f indicator (Badura et al., 2014). (A) 3D plot of network model and microscope configuration. (B) Spike raster for the simulated experiment, where each ROI receives a number of laser pulses equal to its 1-based index. (C) Δ*F/F*_0_ traces for each ROI, showing stronger responses for neurons having spiked more, but varying with expression levels. Heterogeneity in noise is due to varying distances from the focal plane.

**Figure S6:**
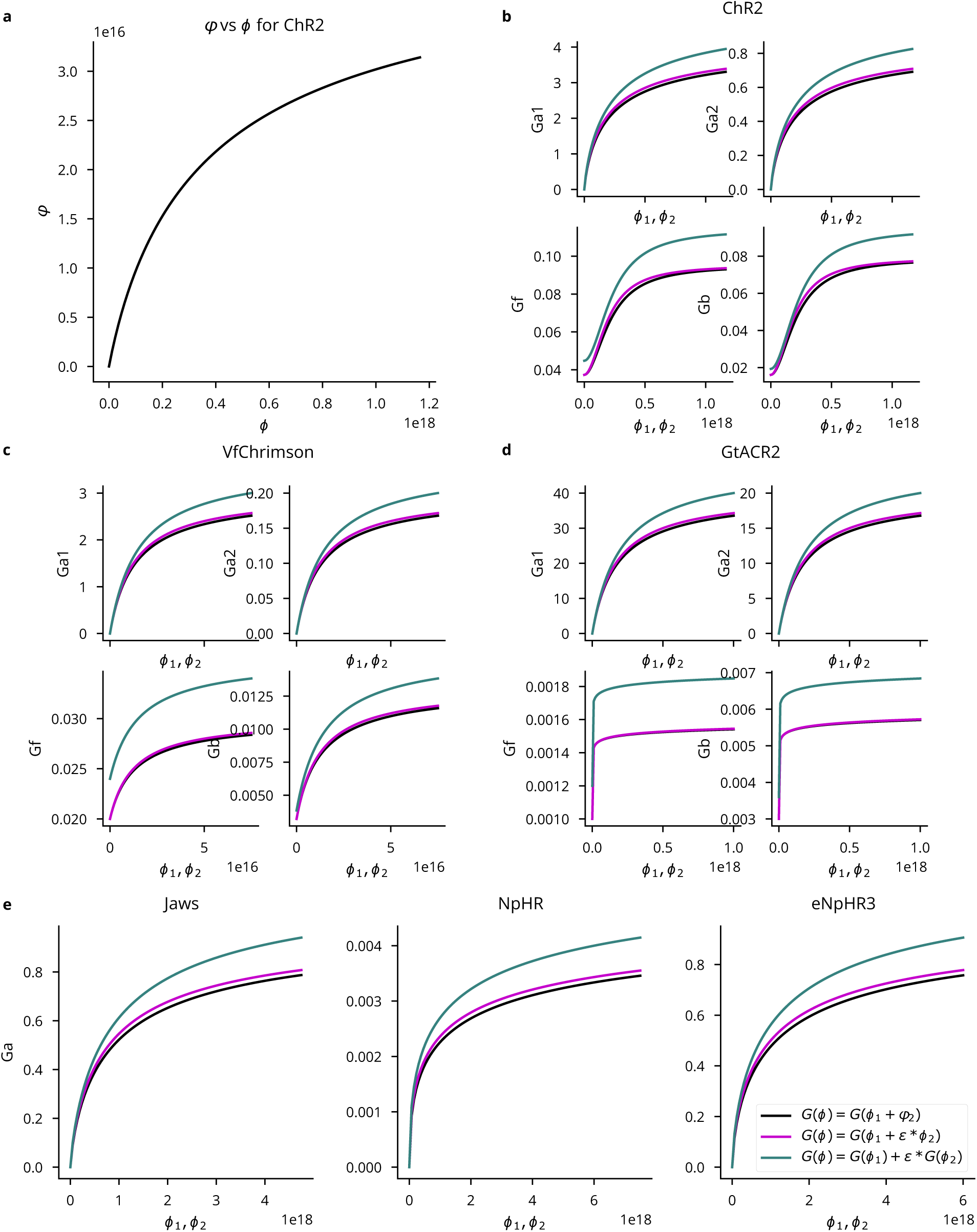
Multi-wavelength opsin model comparison. *ϕ*_1_*, ϕ*_2_ refer to photon flux at peak wavelength *λ*_1_ and some other wavelength *λ*_2_, respectively. All panels take *ε* = 0.2 and use the legend in *E*. (A) The computed effective flux *φ* at *λ*_2_ as a function of the actual flux *ϕ*. (B-D) Light-dependent activation functions for four-state ChR2, Vf-Chrimson, and GtACR2 opsins. (E) Light-dependent activation for the three-state anion pump models. Parameters given in (Bansal et al., 2020b).

**Figure S7:**
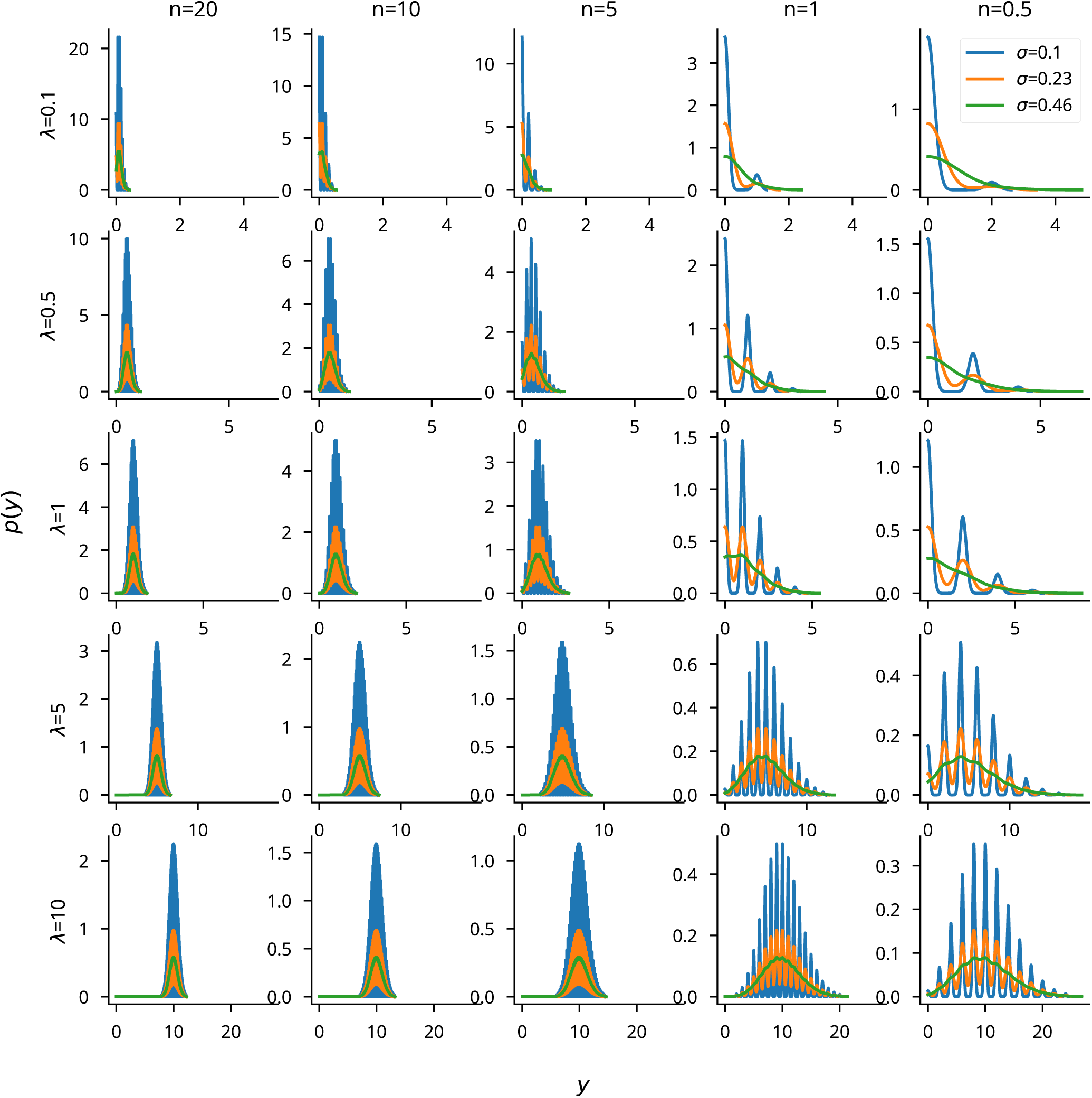
A visualization to assess the appropriateness of the Gaussian noise model for imaging experiments. We plot the Gaussian distribution *p*(*y*) = N (*x, σ*) over a Poisson photon count per pixel *x* ∼ Pois(*λ*). *N* refers to the number of pixels visible in the ROI and *λ* is the expected photon count. Plots show a roughly Gaussian-distributed *p*(*y*) when *N >* 1, which is a realistic assumption for imaging experiments. The spikiness would be mitigated in a real experiment, where *λ* and *σ* would not be constant across pixels. The Gaussian observation appears to be least appropriate for low photon counts, where the distribution has a heavy right tail.

**Figure S8:**
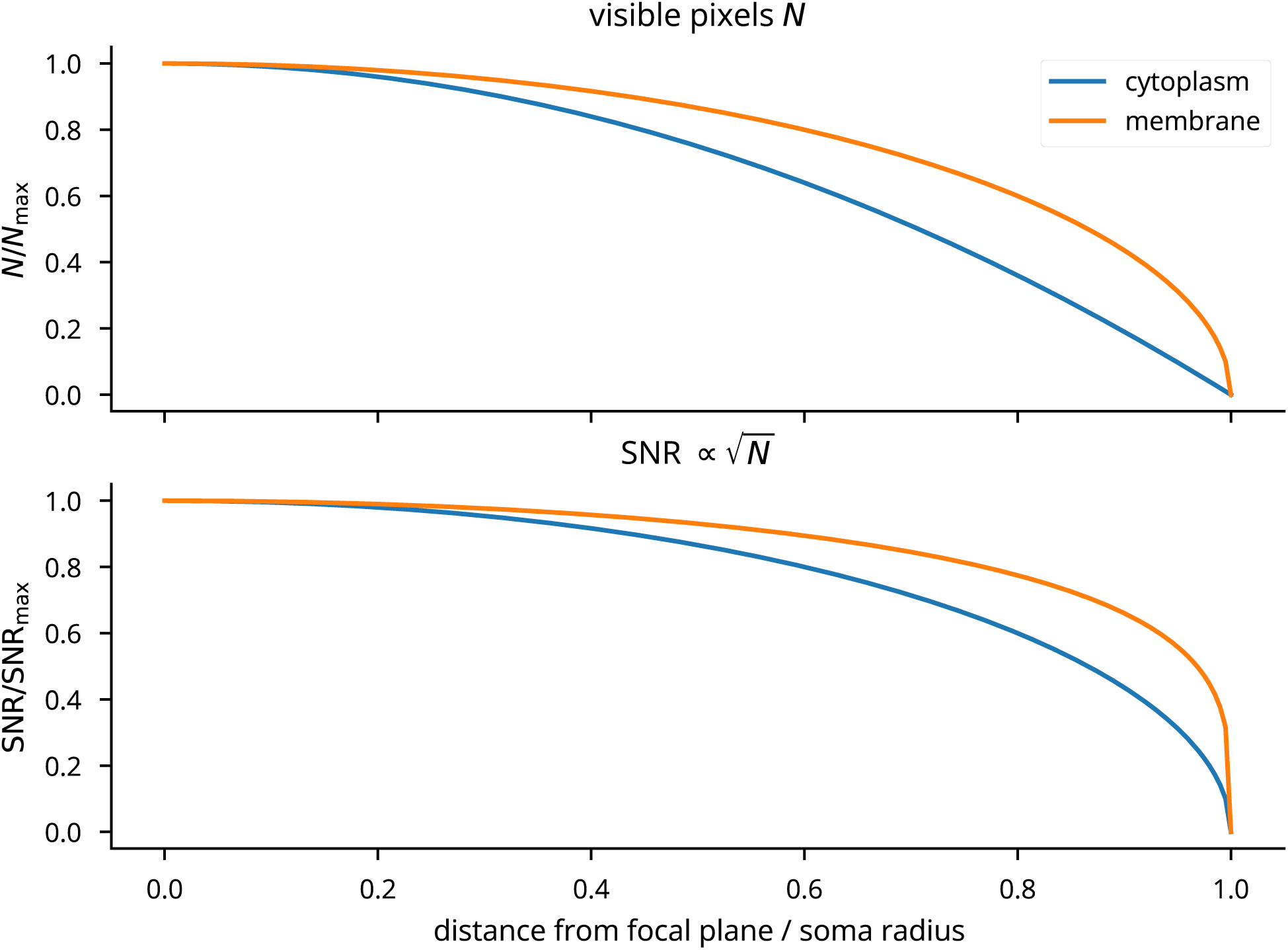
Plot of the number of visible pixels *N* and the SNR as a function of the distance from the focal plane, for indicators found both in the cytoplasm (calcium indicators) and membrane (voltage indicators).

**Figure S9:**
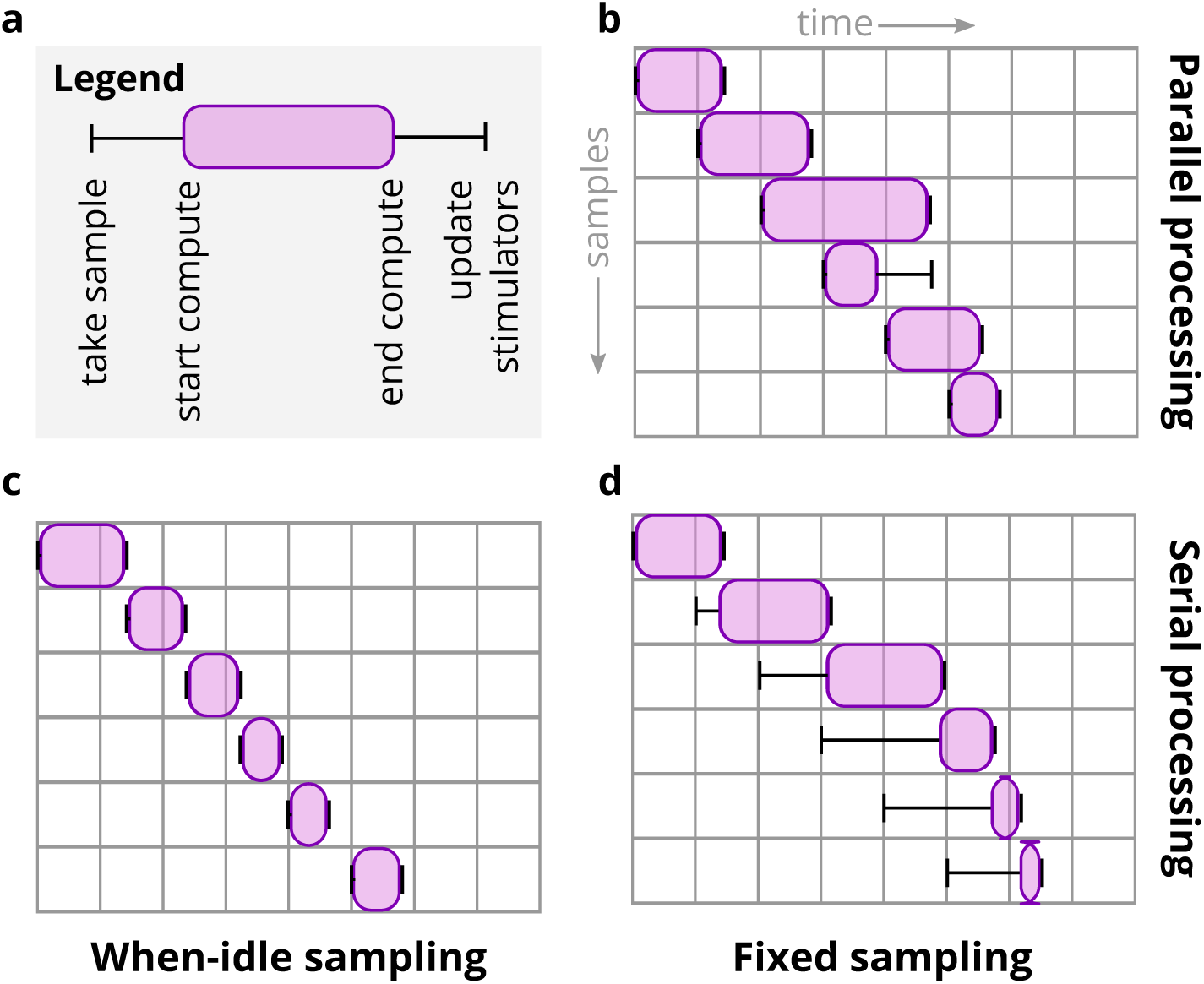
Latency emulation strategy and available configurations. (A) Cleo registers the time a sample is taken from the recording devices, determines the times the computation starts and ends, applies the user-specified delay, and updates stimulation devices when finished. (B) The default parallel processing/fixed sampling mode. Updates are reserved until the previous update is delivered so the sequence of stimulator updates corresponds to the sequence of measurements. (C) The “when-idle” processing mode samples only once the computation for the previous step has terminated. (D) The serial processing/fixed sampling case reflects when computations are not performed in parallel, but sampling continues on a fixed schedule. Samples are taken either as soon as possible after the previous sample time was missed, or on schedule otherwise.

**Figure S10:**
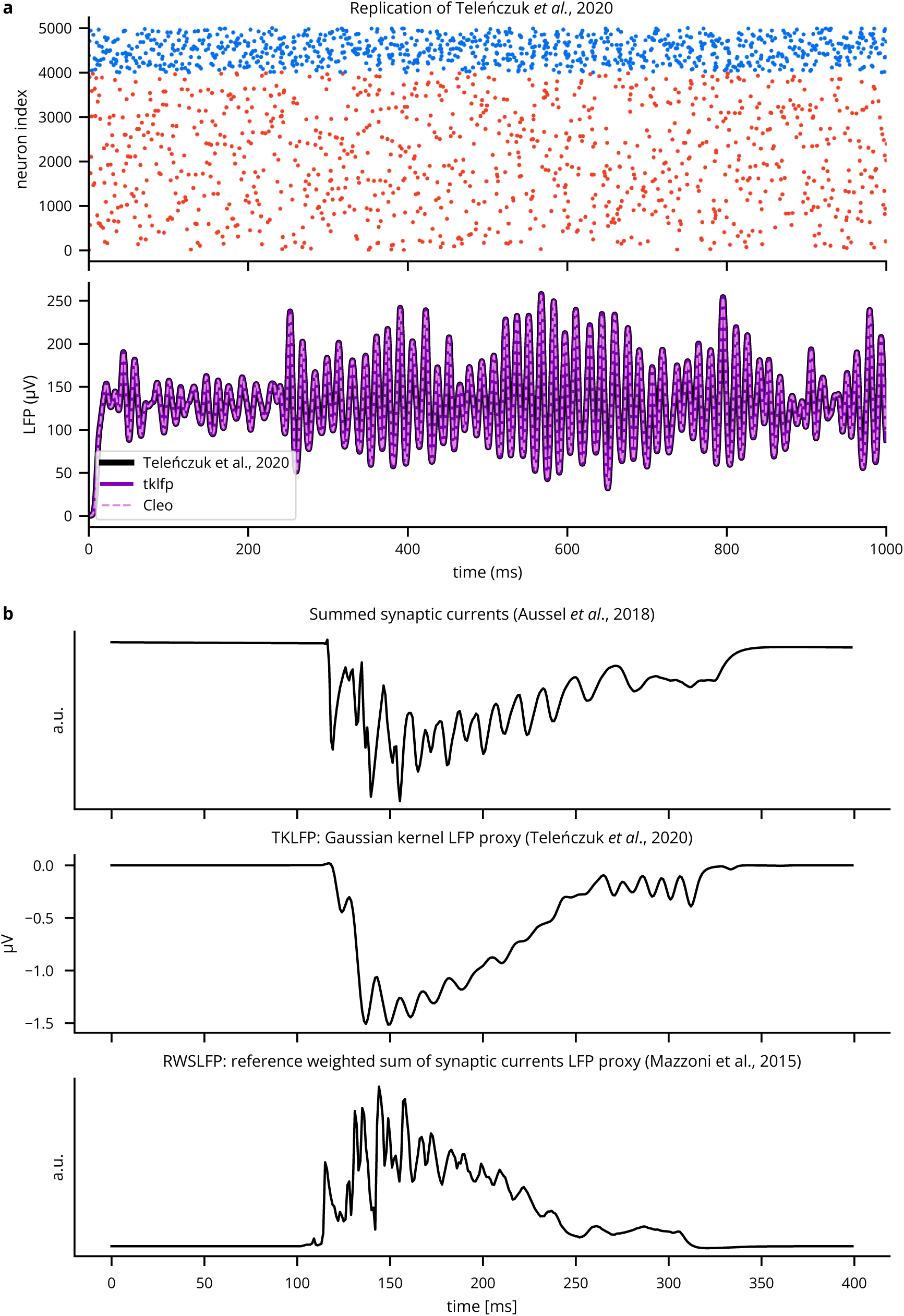
Verification of LFP proxy methods. (A) Replication of the Teleńczuk kernel LFP demo (Telenczuk et al., 2020). (B) Comparison of LFP proxy signals during SWR-like activity in a hippocampus model (see Sec. 3.3.3). Aussel et al. (2018) represent LFP with a sum of synaptic currents, each neuron’s contribution depending on its location in space. The Gaussian kernel approximation method is as described by Telenczuk et al. (2020) and computed by Cleo, which uses the tklfp package implementation (Johnsen, 2022). The reference weighted sum method is described by Mazzoni et al. (2015) and is also computed by Cleo, which uses the wslfp package implementation (Johnsen et al., 2024) (see Sec. 2.4.2).

**Figure S11:**
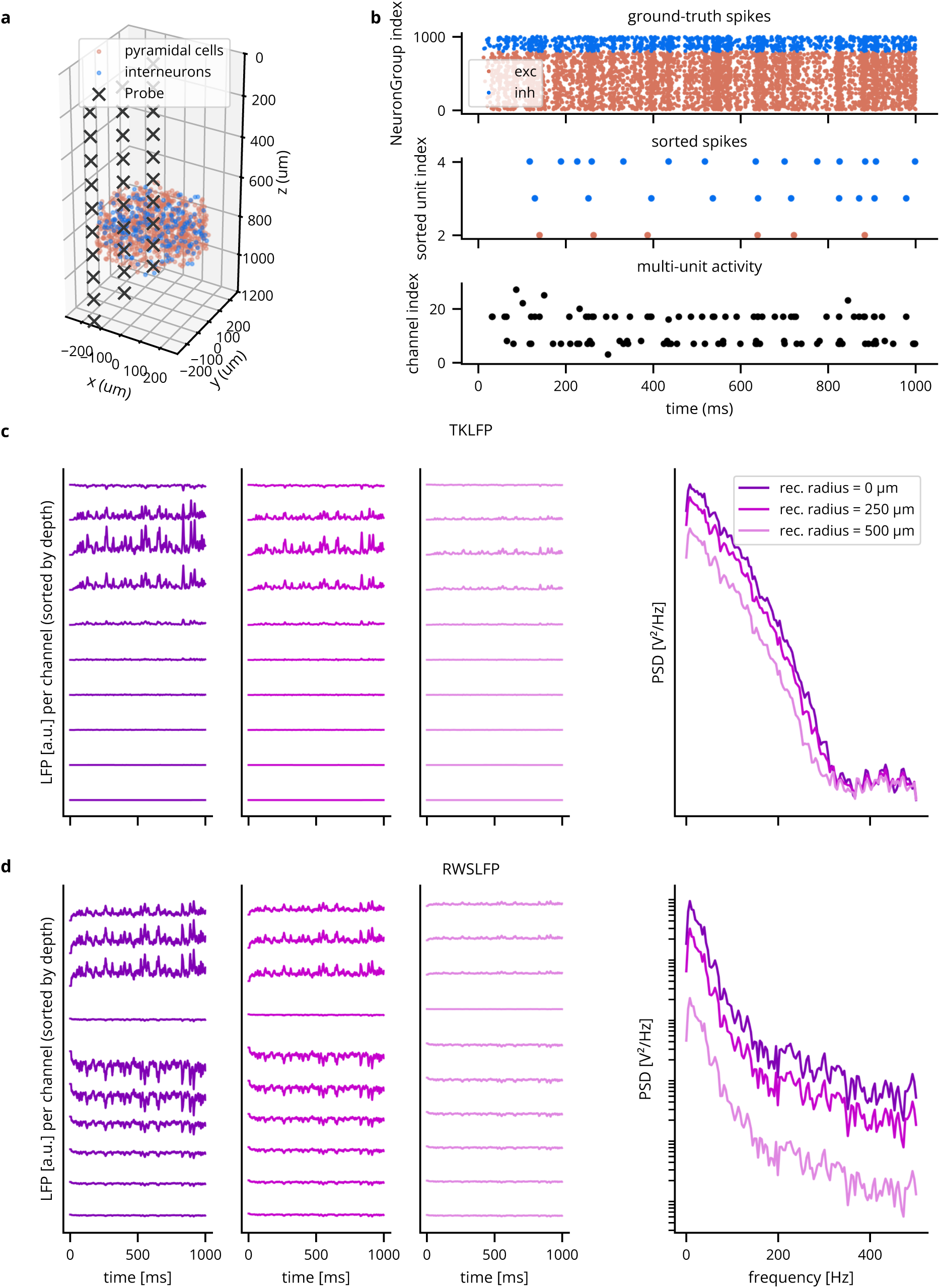
Comparison of the TKLFP and RWSLFP proxy methods for a simulated E/I network. We see here that TKLFP captures less high-frequency content, which is as reported by Teleńczuk *et al*. (A) A Cleo-generated plot of the network model and electrode placement. (B) Sorted and multi-unit spiking activity recorded from the network. (C) LFP and power spectral density (PSD) for the TKLFP signal recorded by the electrod_5_e._1_ (D) Same as *C*, but for the RWSLFP signal.

